# Chemical genetic interaction linking eIF5A hypusination and mitochondrial integrity

**DOI:** 10.1101/2023.12.20.571781

**Authors:** Ken Matsumoto, Rumi Kurokawa, Megumi Takase, Tilman Schneider-Poetsch, Feng Ling, Takehiro Suzuki, Peixun Han, Taisei Wakigawa, Masato Suzuki, Mohammad Tariq, Akihiro Ito, Kyohei Higashi, Shintaro Iwasaki, Naoshi Dohmae, Minoru Yoshida

## Abstract

The eukaryotic translation factor eIF5A plays an important role in translation elongation, especially across stretches of prolines and charged amino acids, and in translation termination. eIF5A undergoes hypusination, a post-translational modification unique to this protein, although the role of hypusination in the function of eIF5A remains elusive. Here, we investigated the cellular defects induced by the hypusination inhibitor GC7 (N1-guanyl-1,7-diaminoheptane). Proteome, translatome, and transcriptome analyses indicated that GC7 downregulated a subset of mitochondrial proteins and DNA, causing mitochondrial stress and eliciting the integrated stress response. Chemical genomic screening using barcoded shRNA libraries identified genes encoding proteins involved in polyamine metabolism/transport and *MPV17L2*, a mitochondrial disease gene homologue whose product regulates mitochondrial translation. Depletion of MPV17L2 caused hypersensitivity to GC7 and phenocopied the effects of GC7 treatment. These results suggest that eIF5A hypusination and MPV17L2 synthetically regulate mitochondrial molecular synthesis and integrity.

---

Translation of cytoplasmic mRNAs is affected by various factors including the nucleotide sequences of mRNAs in the coding region, the combination of cis-acting elements in untranslated regions and cognate trans-acting factors, nuclear and cytoplasmic history of mRNAs that influence the architecture of messenger ribonucleoproteins, availability of amino acids and tRNAs, and post-translational modifications of translation factors.^1–5^ Stalling of a ribosome during translation elongation causes the trailing ribosome(s) moving on the same mRNA to collide.^6,7^ The eukaryotic initiation factor 5A (eIF5A) is involved in resolving ribosome slowdown due to slow peptidyl transfer in the cytoplasm,^8,9^ thereby supporting mitochondrial function.^10^ ^11,12^

eIF5A was originally identified as an initiation factor that facilitates the first peptide bond formation, as determined by the methionyl (Met)-puromycin synthesis assay; however, later studies showed that it functions mainly during translation elongation and termination.^13,14^ eIF5A binds near the E site of ribosomes and promotes the stabilization of the peptidyl tRNA at the P-site, which facilitates efficient peptide bond formation, especially at the Pro-Pro peptide bond^15^ or at non-Pro-Pro peptide bonds more broadly.^9,16^

eIF5A contains the unique amino acid hypusine.^17–19^ The formation of a hypusine residue, hypusination, occurs at a conserved and specific lysine residue in eIF5A (Lys50 in human eIF5A). The hypusine moiety of eIF5A interacts with the CCA-end of the peptidyl tRNA at the peptidyl transferase center of the ribosome.^15,20^ The amino acid at the structurally conserved site of EF-P, the bacterial ortholog of eIF5A, is also post-translationally modified with β-lysine or rhamnose,^21–23^ whereas that of archaeal IF5A is hypusinated.^24^ The conservation of hypusine and equivalent modifications across three domains of life underscores the significance of hypusine modification for eIF5A function.

Hypusination is catalyzed by a two-step enzymatic cascade. An aminobutyl moiety of spermidine, one of three polyamines in animal cells, is transferred to the lysine residue by deoxyhypusine synthase (DHS or DHPS), converting it to deoxyhypusine.^25,26^ Deoxyhypusine is subsequently hydroxylated by deoxyhypusine hydroxylase (DOHH) to form mature hypusine.^27,28^ The (deoxy)hypusination is essential for the function of eIF5A in Met-puromycin formation,^14,29^ binding to ribosomes, translation elongation and termination, ^9,29,30^ and start codon selection.^31,32^ Hypusinated eIF5A plays roles in cell proliferation,^33^ autophagy,^34,35^ macrophage activation,^12^ anti-microbial responses in macrophages,^36^ helper T cell lineage fidelity,^37^ and brain aging and memory.^38,39^

Because eIF5A, DHS, and DOHH are all essential for cell proliferation in higher eukaryotes,^40,41^ compounds targeting the hypusination reaction provide a useful tool. A search for small compounds capable of inhibiting the *in vitro* DHS reaction identified N1-guanyl-1,7-diaminoheptane (GC7).^42^ GC7 is structurally similar to spermidine, and thus competes with spermidine for DHS binding.^43^ GC7 has been widely used as a hypusination inhibitor in cultured cells. GC7 has different effects on cells, including anti-proliferation,^33,44^ anti-tumor,^44–47^ and anti-apoptosis effects,^11,44^ promoting mitochondrial dysfunction,^11,12^ and inhibiting translation initiation and elongation.^48^ However, the mechanism by which hypusination deficiency induces these phenotypic changes remains elusive.

Here, we performed genetic screening to search for genes that alter the cellular sensitivity to GC7. The results show that GC7 inhibits growth by acting on the polyamine pathway, especially polyamine biosynthesis and eIF5A hypusination in cells. We identified *MPV17L2* as a novel synthetic lethal gene that is critical for maintaining cell viability in the presence of GC7. MPV17L2 was required for robust mitochondrial translation and mitochondrial DNA (mtDNA) maintenance. The present results indicate that hypusinated eIF5A is crucial for the maintenance of the two protein synthesis systems in the cytoplasm and mitochondria.

## Results

### GC7 treatment downregulates mitochondrial proteins

To explore the effect of hypusination inhibition on the cellular proteome, we performed proteomic analysis of HeLa cells treated with GC7 for 24 h and 72 h (Fig. 1A–1D and S1). LC-MS/MS identified 6,070 proteins based on quantitative data, of which 5,390 proteins were detected in both 24 and 72 h treated cells. The number of proteins downregulated (<0.5-fold) by GC7 was higher than that of proteins upregulated (>2-fold) by GC7 at both time points, which is consistent with previous reports that the inhibition of hypusination by GC7 inhibits translation at both initiation and elongation steps.^48^ Pathway analysis of proteins downregulated >2-fold by GC7 treatment for 24 h showed that those particularly affected were proteins involved in oxidative phosphorylation (p = 1.6 × 10^-35^) and mitochondrial dysfunction (p = 1.0 × 10^-34^) (Fig. 1A–1F). Among the 656 mitochondrial proteins identified in the proteome analysis, 160 and 226 proteins showed >2-fold downregulation after GC7 treatment for 24 h and 72 h, respectively, whereas only 7 and 23 proteins were upregulated >2-fold. These results are consistent with previous findings,^11,12^ and confirm that GC7 treatment downregulates a subset of mitochondrial proteins including components of the respiratory chain.

**Figure 1.**
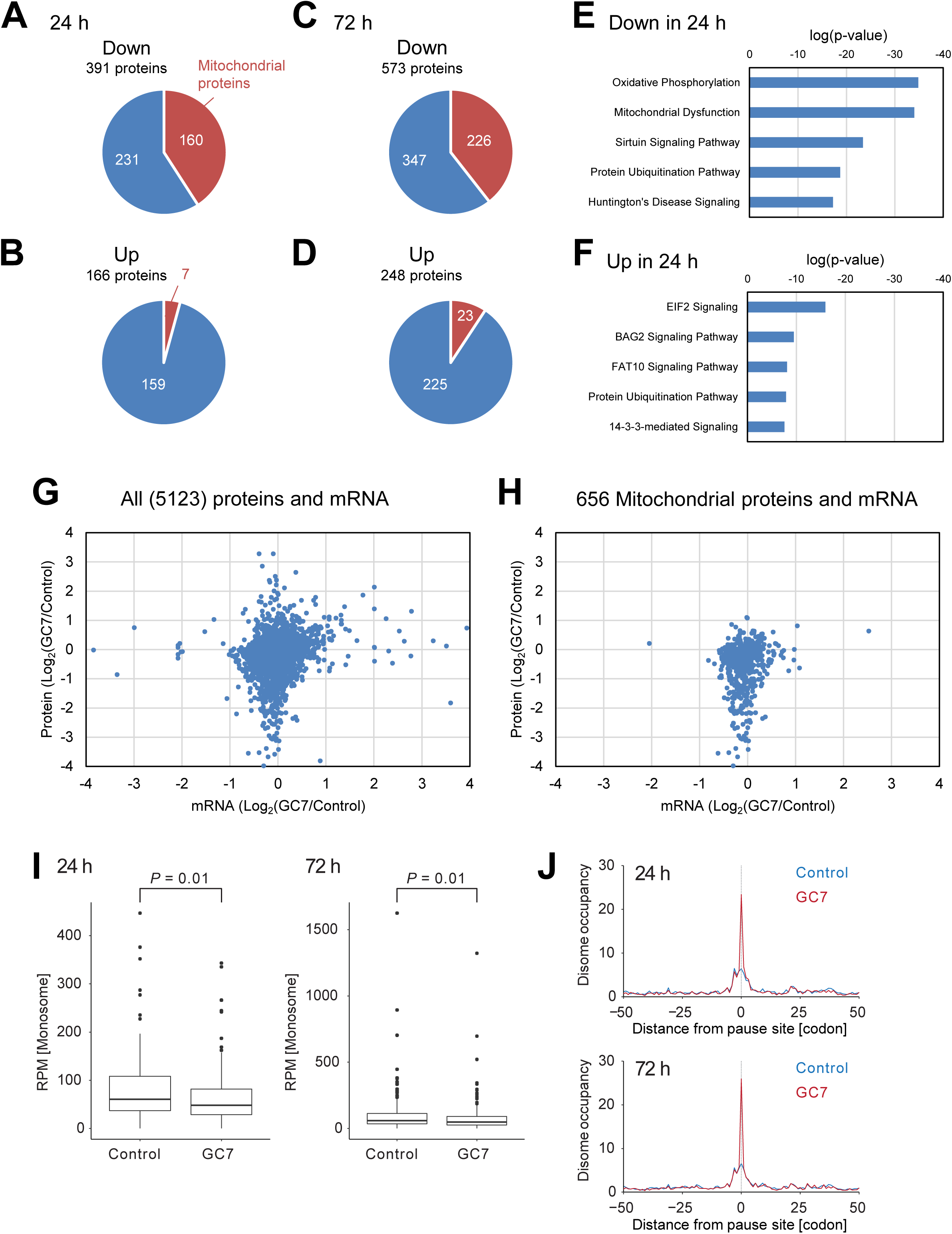
GC7 treatment downregulates mitochondrial proteins at the post-transcriptional level. (A-F) Proteome analysis of GC7-treated cells. HeLa S3 cells were treated with 10 μM GC7 for 24 h or 72 h. Cellular proteins were analyzed by label-free mass spectrometry. Proteins downregulated (<0.5-fold, A and C) or upregulated (>2-fold, B and D) in cells treated for 24 h (A and B) or 72 h (C and D) compared with untreated cells were analyzed against the list of mitochondrial proteins from MitoCarta 3.0. Proteins downregulated (E) or upregulated (F) in cells treated with GC7 for 24 h were subjected to pathway analysis. (G and H) Effect of GC7 on the abundance of mRNA and protein. The abundance of commonly detected mRNAs from RNA-seq data and proteins from proteome data of cells treated with 10 μM GC7 for 24 h was compared (G). The abundance of mitochondrial proteins and their encoding mRNAs was compared (H). (I) Box plot of ribosome footprint reads of mitochondrial genes under the indicated conditions. In the box plots, the median (center line), upper/lower quartiles (box limits), and 1.5× interquartile range (whiskers) are shown. RPM, reads per million mapped reads. (J) Metagene plots of disome occupancy around the disome pause sites under the indicated conditions.

To determine whether mitochondrial protein downregulation by GC7 treatment was associated with changes in the transcriptome, we performed RNA-seq analysis (Fig. S1). We compared the changes in the levels of 5,123 transcripts/proteins commonly detected in transcriptome and proteome analyses (Fig. 1G). The results showed that protein levels decreased more substantially than transcript levels, with a small overlap between the changes in transcripts and proteins. This trend was also observed in the analysis of mitochondrial proteins and suggests that the downregulation of mitochondrial proteins occurs mainly at the post-transcriptional level (Fig. 1H).

To further investigate the effect of GC7 on translational control, we performed ribosome profiling, a deep sequencing-based method for analyzing ribosome-protected RNA fragments following RNase treatment.^49,50^ Consistent with the data above, we detected reduced footprint reads from genes encoding mitochondrial proteins, which were downregulated by GC7 treatment in the proteome analysis (Fig. 1I).

Because eIF5A facilitates translation elongation,^9,15,16,51^ we hypothesized that GC7 may induce ribosome stalling and subsequent collision with the trailing ribosome to form di-ribosomes or disomes. To test this, GC7-treated cells were analyzed by disome profiling, which sequences longer footprints generated by collided ribosomes.^8,52–54^ The analysis detected the more frequent ribosome collision that occurred in the mRNAs encoding mitochondrial proteins, which were downregulated by GC7 treatment in the proteome analysis (Fig. 1J).

Taken together, the data suggest that hypusination of eIF5A ensures the efficient synthesis of mitochondria-localized proteins by preventing ribosome stalling on the associated mRNAs.

### GC7 treatment decreases mitochondrial protein and DNA levels

The results obtained led us to investigate mitochondrial integrity in the presence of GC7. GC7 treatment markedly decreased the levels of various mitochondrial proteins, including components of respiratory chain complexes (OXPHOS) I–IV, TFAM, HADHA, mitochondrial RNA polymerase (POLRMT), and the mitochondrial ribosomal protein MPRS17 (Fig. 2A and 2B). These results were consistent with those of proteome analysis, although the effects on ATP5A and ATP5B, which are subunits of FoF1-ATP synthase (or complex V), were less pronounced. Alterations in mRNA abundance assessed by qRT-PCR did not explain the changes in ATP5A, UQCRC2, SDHB, and NDUFB8 protein levels (Fig. 2C).

**Figure 2.**
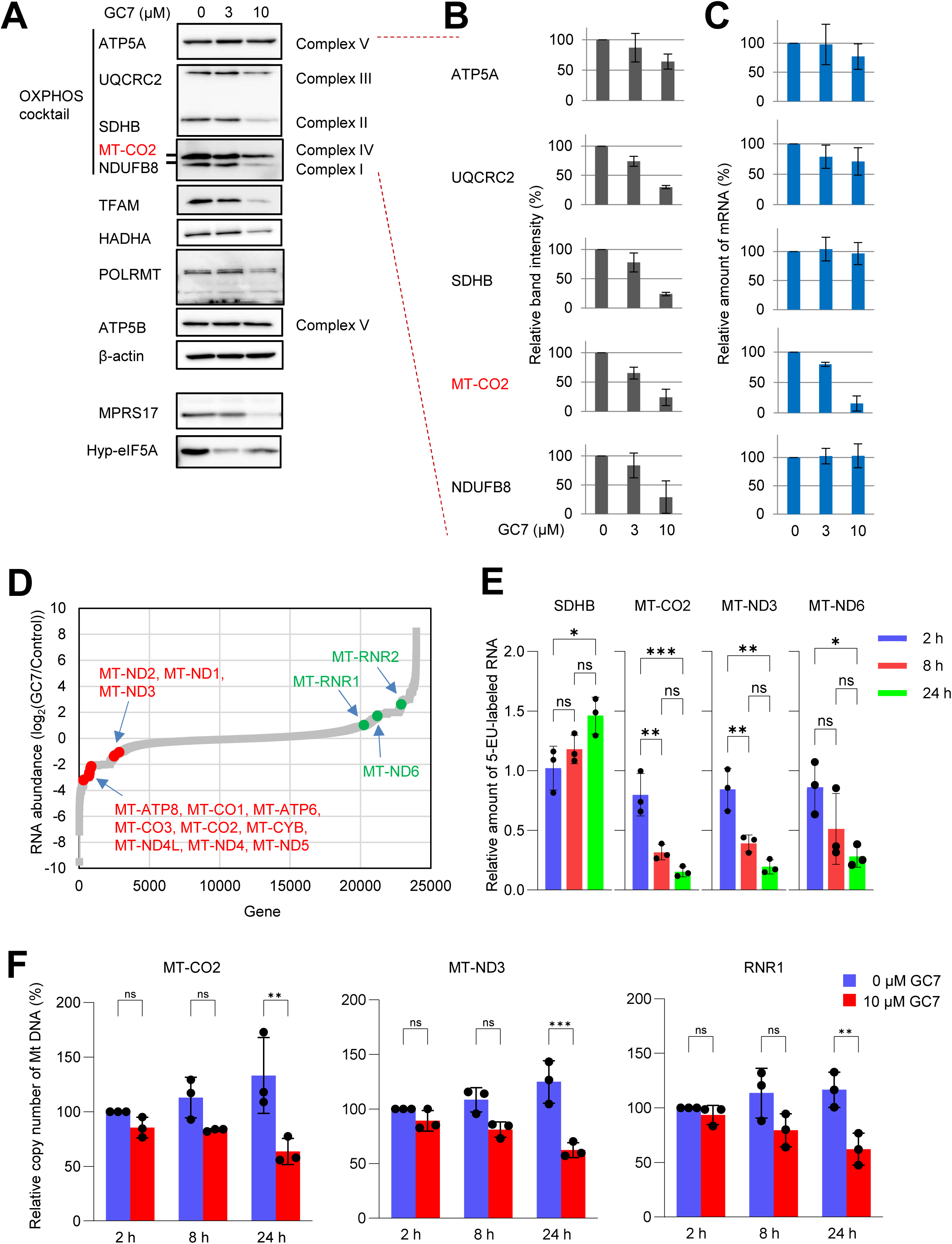
Decrease in mitochondrial transcription and the levels of mitochondrial genome DNA induced by GC7 treatment. (A-C) HeLa S3 cells were treated with GC7 for 24 h, and cellular proteins were analyzed by western blotting (A). The bands corresponding to five OXPHOS proteins were quantified (B). The corresponding mRNAs from the cells were analyzed by qPCR (C). Among the five mRNAs, only MT-CO2 (indicated in red) is encoded by mtDNA. (D) RNA-seq analysis of GC7-treated cells. RNA abundance was compared between untreated and GC7-treated cells. RNAs encoded by mtDNA are indicated. (E) Mitochondrial transcriptional activity was examined. Cells treated with GC7 were pulse-labeled with EU, and the EU-labeled RNAs were analyzed for nuclear-encoded (SDHB) and three mitochondrially-encoded mRNAs by RT-qPCR. (F) The amount of mtDNA in cells treated with 10 μM GC7 was analyzed by qPCR. Data are expressed as the mean ± S.D. \**P* < 0.05, \*\**P* < 0.01, \*\*\**P* < 0.001, \*\*\*\**P* < 0.0001 (One-way ANOVA).

In contrast to these genes, the mRNA encoding MT-CO2, a gene in mtDNA, was decreased by GC7 treatment (Fig. 2C). Considering that mtDNA encodes 13 mRNAs corresponding to subunits of complexes I, III, IV, and V, we analyzed the RNA-seq data from GC7-treated cells and found that 12 mtDNA-encoded mRNAs were significantly decreased (Fig. 2D). MT-ND6 mRNA and two mitochondrial ribosomal RNAs (MT-RNR1 and MT-RNR2) lacking poly(A) tails,^55,56^ were not downregulated by GC7, which could be attributed to poly(A)-selection in the RNA-seq procedure. The downregulation of the 13 mtDNA-encoded mRNAs and 2 rRNAs was confirmed by qRT-PCR (Fig. S2).

Next, we examined the transcriptional activity of mtDNA-encoded mRNAs. Cells were treated with 10 μM GC7 for 2, 8, and 24 h, and nascent transcripts were labeled with 5-ethynyl uridine (EU) during the final 1 h of incubation before harvesting (Fig. 2E). The EU-labeled RNA was captured from total cellular RNA using a click reaction and analyzed by qRT-PCR. All mtDNA-encoded mRNAs tested in this assay were downregulated by GC7 in a time-dependent manner, reaching 20–30% of the control after 24 h of treatment (Fig. 2E). This indicated that GC7 inhibited transcription in mitochondria. GC7 had no effect on the transcription rate of the nuclear-encoded SDHB mRNA (Fig. 2E).

To further explore the decrease in mRNA synthesis in mitochondria, the mtDNA content in GC7-treated cells was examined by qPCR (Fig. 2F). GC7 treatment for 24 h decreased the amount of mtDNA to 60% of the control. Although GC7 treatment decreased both, mitochondrial DNA and RNA, the decrease in mitochondrial RNA was more pronounced than that of DNA.

These results suggest that mitochondrial proteome downregulation by GC7 involves two different mechanisms: the amounts of nuclear-encoded proteins are regulated at the post-transcriptional level, whereas the downregulation of mtDNA-encoded proteins is a result of the decrease in mRNA levels. Nuclear-encoded OXPHOS subunits are imported in large excess into mitochondria, indicating that mitochondrial translation is the rate-limiting step for OXPHOS assembly, and unassembled subunits are rapidly degraded.^57,58^ The present results may reflect the decrease in the synthesis of mtDNA-encoded OXPHOS subunits resulting from loss of mtDNA and mRNA inside the organelle.

### Changes in mitochondrial morphology caused by GC7 treatment

Mitochondria are dynamic organelles that continuously undergo fission and fusion. These processes can be hampered by mitochondrial stress inducers, resulting in morphological changes such as fragmentation or hyperfusion.^59,60^ Because GC7 affected the levels of mitochondrial DNA, RNA, and proteins, we examined mitochondrial morphology by staining with Mitotracker Red CMXRos, a membrane potential-dependent dye, and immunofluorescence analysis with an antibody against Tom20, a mitochondrial outer membrane protein. In GC7-treated cells, anti-Tom20 staining detected mitochondrial fragmentation, and doughnut-like structures showing faint Mitotracker Red fluorescence were observed (Fig. 3A). The doughnut-like structures were observed after treatment with the mitochondrial oxidative phosphorylation uncoupler FCCP and the FoF1 ATP synthase inhibitor oligomycin A, as reported previously,^61,62^ suggesting that these structures were associated with the attenuated membrane potential. In GC7-treated cells, we also detected small punctate structures strongly emitting Mitotracker Red fluorescence that did not co-localize with Tom20, although the origin of these signals remains unclear.

**Figure 3.**
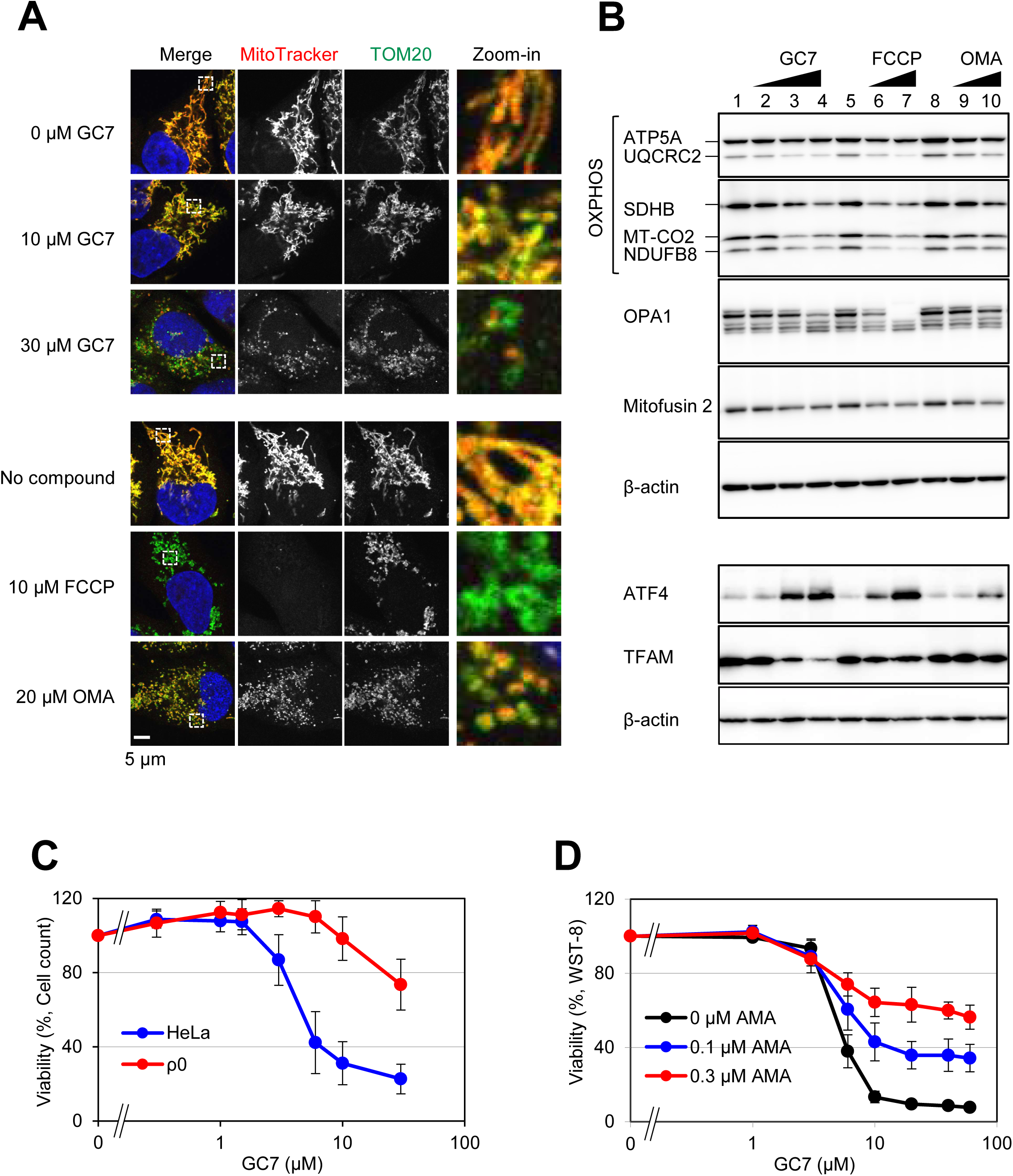
GC7 induces mitochondrial fragmentation. (A and B) HeLa S3 cells were treated with GC7, FCCP, and oligomycin A (OMA) for 24 h. Mitochondrial morphology was analyzed by staining with Mitotracker Red followed by immunofluorescence detection with anti-Tom20 antibodies. The cells were counter-stained with DAPI and observed by confocal microscopy (A). Cell lysates were analyzed by western blotting (B). (C) HeLa cells and HeLa ρ0 cells were treated with different concentrations of GC7 for 72 h and counted. (D) The GC7 sensitivity of HeLa S3 cells co-treated with antimycin A (AMA) was analyzed using the WST-8 assay.

Proteolytic processing of optic atrophy 1 (OPA1), a mitochondrial inner membrane dynamin-like GTPase, is linked to mitochondrial fragmentation, whereas the outer membrane GTPases Mitofusin 1 and 2 are involved in mitochondrial fusion. To test the involvement of these proteins in the aberrant mitochondrial shape in GC7-treated cells, we performed western blot analysis. GC7 induced processing of OPA1 and slightly downregulated Mitofusin 2, consistent with the change in mitochondrial morphology (Fig. 3B). A similar phenotype was observed in the presence of FCCP or oligomycin A treatment. These results indicate that GC7 causes mitochondrial fragmentation.

The observed effects of GC7 indicated that it is not a direct uncoupler of membrane potential. Unlike FCCP or oligomycin A, which cause mitochondrial fragmentation after incubation periods as short as 1 h,^62^ at least 3 or 4 h of GC7 treatment were required for the induction of mitochondrial fragmentation (Fig. S2).

### GC7 activates the integrated stress response through mitochondrial stress

Mitochondrial stress inducers such as actinonin, doxycycline, FCCP, and MitoBloCK-6 activate the integrated stress response (ISR), thereby upregulating ATF4 transcription and translation.^63^ We examined the levels of ATF4 in GC7-treated cells, which showed that GC7 upregulated the ATF4 protein in a dose-dependent manner (Fig. 3B). Taken together with previous data showing that GC7 induces eIF2α phosphorylation and inhibits global translation initiation,^48^ and that eIF5A depletion induces the ER stress markers IRE-1α, phosphorylated PERK, and ATF4,^64^ the present results indicate that GC7 activates the ISR by inducing mitochondrial stress.

To further examine the role of mitochondria in the cytotoxicity of GC7, we tested the GC7 sensitivity of ρ0 cells, which are devoid of mtDNA (Fig. 3C). ρ0 cells derived from HeLa cells were less sensitive to GC7 than parental HeLa cells (Fig. 3C), demonstrating that mitochondria play a role in GC7 cytotoxicity. Combination treatment with antimycin A, an inhibitor of the respiratory chain complex III, decreased the sensitivity of cells to GC7 (Fig. 3D). These results indicate that GC7 acts as a stress inducer in mitochondria and its cytotoxicity depends on mitochondrial activity.

### shRNA screening for genes that determine GC7 sensitivity

To identify genes involved in GC7-mediated cytotoxicity, we performed genome-wide shRNA screening.^65^ Approximately 15,000 human genes were targeted in separate transductions using DECIPHER lentiviral library Human Modules 1, 2, and 3, each of which contains 27,500 shRNAs against approximately 5,000 independent genes (five or six shRNAs per gene). HeLa S3 cells transduced with the libraries were treated with or without 10 μM GC7 for 10 days (Experiment 1, Exp. 1) or 14 days (Exp. 2) (Fig. 4A). The shRNA-specific barcodes were PCR-amplified from the genomic DNA of GC7-treated and untreated cells and subjected to next-generation sequencing. The normalized read counts of each barcode were compared between treated and untreated cells. shRNA-targeted genes showing significantly increased (candidate resistance genes) or decreased (candidate sensitivity genes) barcode read counts were considered (Fig. 4B).

**Figure 4.**
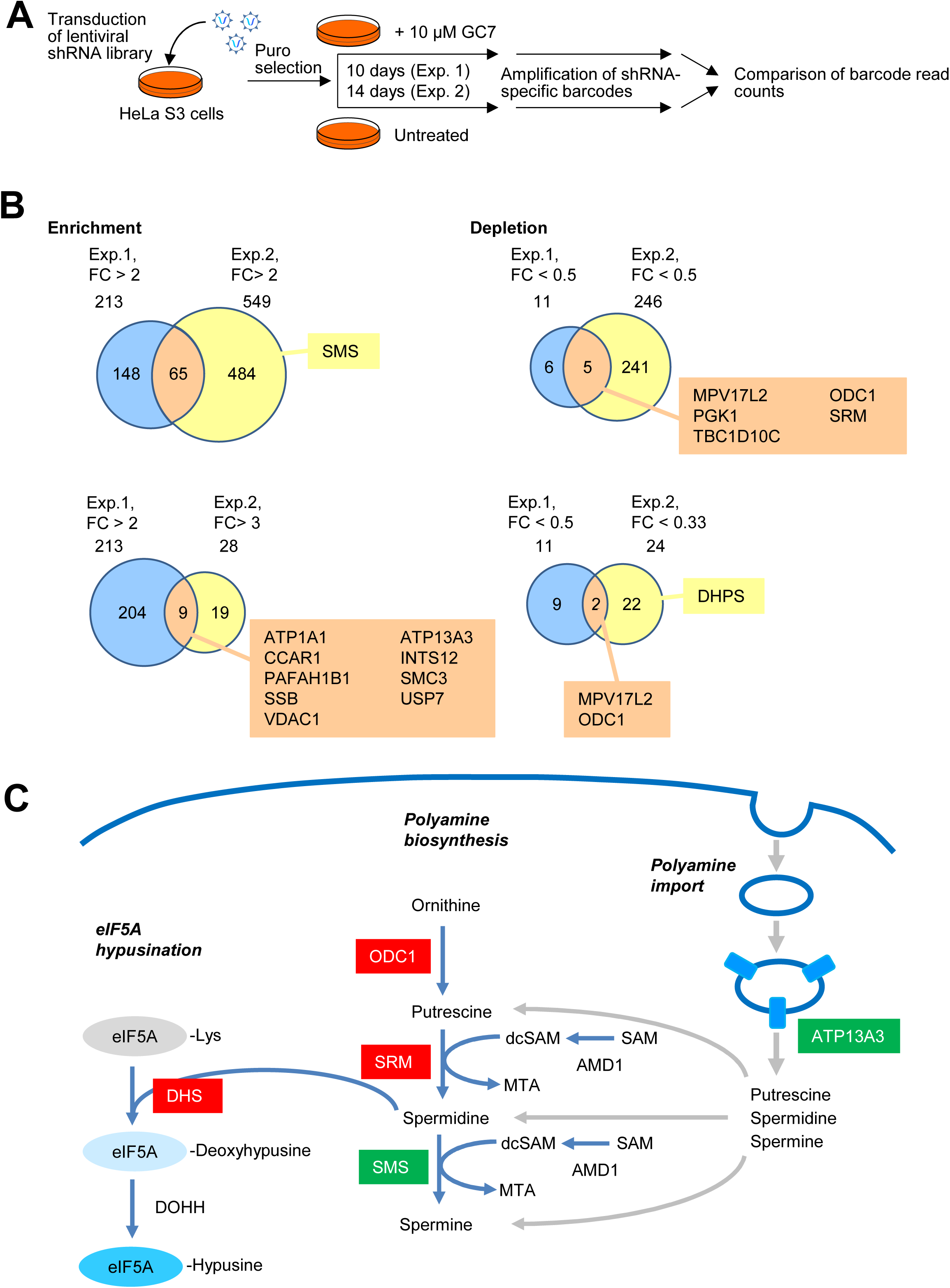
shRNA screening with GC7. (A) Schematic of the shRNA library screening. (B) Comparison of enriched and depleted shRNAs between two independent shRNA screens. FC, fold change (GC7-treated/untreated). (C) shRNAs against genes involved in polyamine biosynthesis, polyamine import, and hypusination were identified in the shRNA screens. Genes corresponding to shRNAs depleted in the screens are shown in red, and genes corresponding to enriched shRNAs are shown in green.

The resistance genes identified in both Exp. 1 and Exp. 2 included *ATP13A3*, which encodes a recently identified polyamine transporter involved in polyamine import by endocytosis.^66,67^ The sensitivity genes included polyamine biosynthesis pathway genes such as ornithine decarboxylase 1 (*ODC1*) and spermidine synthase (*SRM*) (Fig. 4B and 4C). Depletion of ODC1 and SRM leads to impaired synthesis of spermidine, the substrate of DHS; therefore, the high sensitivity to knockdown of these genes could be explained by the hypomodification of eIF5A and suggests that the cytotoxicity of GC7 involves inhibition of eIF5A hypusination (Fig. 4C).

The sensitivity genes included *MPV17L2*, which encodes a mitochondrial protein (Fig. 4B). The six shRNAs targeting MPV17L2 from the shRNA library Module 3 were depleted in GC7-treated samples with fold change (GC7-treated/untreated) <0.5 in both Exp. 1 and Exp. 2 (Fig. 4B and 5A). *MPV17L2* is a homologue of *MPV17*, which is conserved from yeast to humans and is a causal gene of a human mitochondrial disease caused by mtDNA depletion.^68,69^ MPV17L2 facilitates mitochondrial translation by acting as a mitochondrial ribosome assembly factor.^70^ siRNA-mediated knockdown of MPV17L2 inhibits protein synthesis in mitochondria and causes aggregation of mitochondrial nucleoids.^70^ To the best of our knowledge, there are no reports describing the relationship between MPV17L2 and hypusination or polyamines. We therefore focused on MPV17L2 in the context of GC7 sensitivity in subsequent experiments.

**Figure 5.**
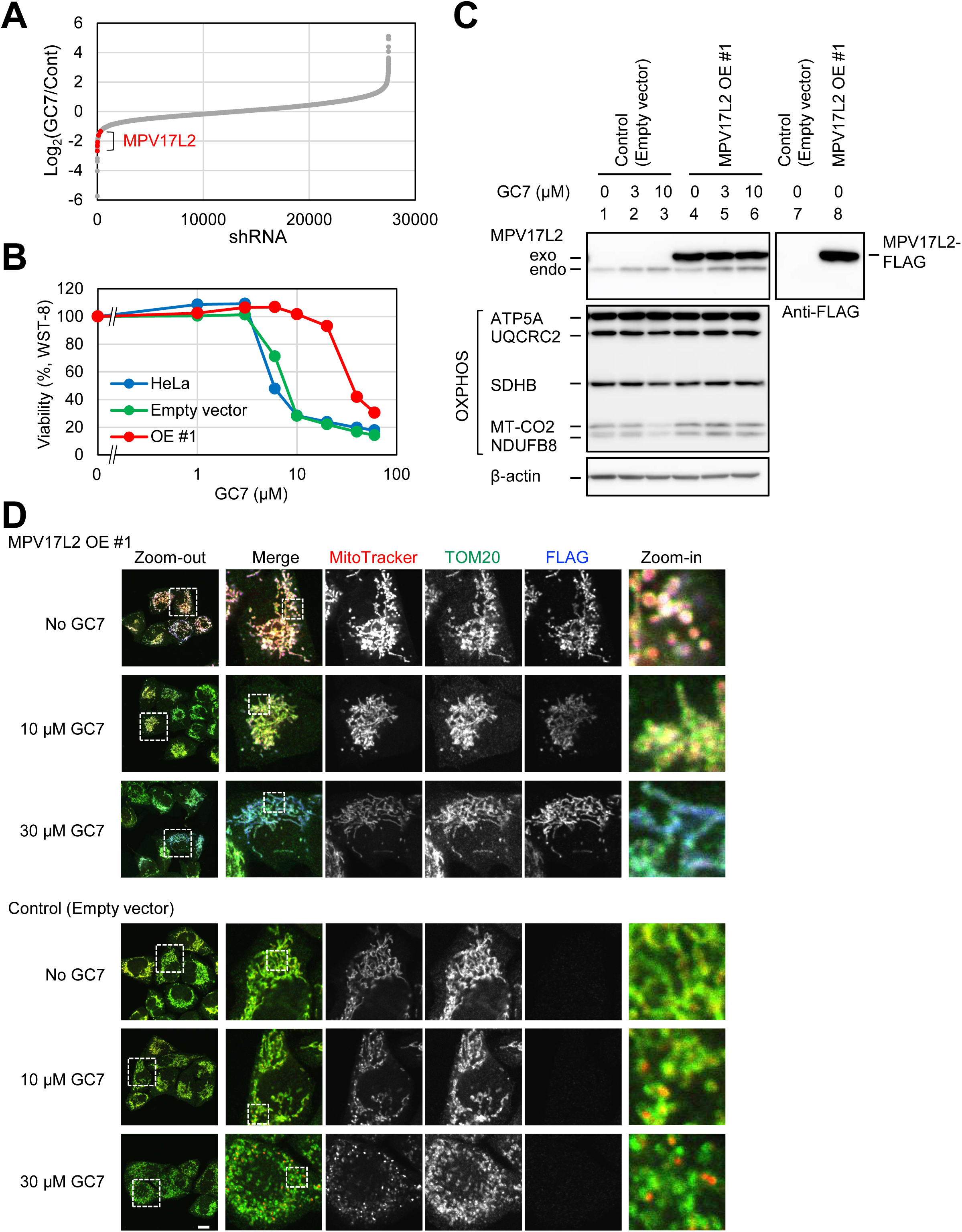
MPV17L2 overexpression decreases the sensitivity of cells to GC7. (A) Fold change of 27,500 shRNAs in the Human Module 3 library in experiment 1. shRNAs for MPV17L2 are shown as red dots. (B) The GC7 sensitivity of parental HeLa S3 cells, empty vector-introduced cells, and MPV17L2-FLAG overexpressing cells (#OE 1) were analyzed by WST-8 assay. (C) #OE 1 cell lysates were analyzed by western blotting. (D) Mitochondrial morphology in #OE 1 cells after GC7 treatment for 24 h was examined by Mitotracker Red staining and immunofluorescence with anti-Tom20 and anti-FLAG antibodies.

### MPV17L2 overexpression restores GC7 cytotoxicity in mitochondria

Given that shRNA-mediated knockdown of MPV17L2 sensitized cells to GC7, we next examined the effect of MPV17L2 overexpression on the sensitivity to GC7. We prepared a stable cell line overexpressing MPV17L2-FLAG in HeLa S3 cells (OE#1 cells, Fig. 5B). OE#1 cells showed decreased sensitivity to GC7 (Fig. 5B), and exposure to GC7 concentrations up to 10 μM for 24 h did not downregulate mitochondrial proteins (Fig. 5C). Immunofluorescence analysis showed that the MPV17L2-FLAG protein localized to mitochondria and colocalized with Tom20 (Fig. 5D), although the mitochondria of OE#1 cells were fragmented and showed strong Mitotracker Red staining, probably due to stress caused by an excess of this protein. OE#1 cells maintained mitochondrial network structures even in the presence of 10 μM and 30 μM GC7 (Fig. 5D). These results indicated that MPV17L2 is a key factor for the maintenance of mitochondrial integrity upon GC7 treatment.

### Depletion of MPV17L2 results in GC7 hypersensitivity and a decrease in mitochondrial DNA

The shRNA screening showed that knockdown of MPV17L2 sensitized cells to GC7 (Fig. 4), and its overexpression decreased sensitivity to GC7 (Fig. 5), suggesting a synthetic lethal genetic interaction between MPV17L2 and DHS. However, the possibility that GC7 targets MPV17L2, thereby showing higher cytotoxic activity, cannot be ruled out. To clarify the mechanism by which MPV17L2-depleted cells are highly sensitive to GC7, we prepared MPV17L2 knockout (KO) cells by CRISPR/Cas genome editing (Fig. 6A). DNA sequencing of the *MPV17L2* gene locus showed that three alleles of the gene in HeLa S3 cells (owing to a karyotypic abnormality) were fully knocked out in KO clone #1 generated using gRNA1 and partially knocked out in clone #2 generated using gRNA2 (Fig. S3). If MPV17L2 is a direct target of GC7 and thus responsible for the cytotoxicity of GC7, *MPV17L2* knockout cells would be resistant to this compound because of the loss of the target protein. However, two *MPV17L2* KO clones were more sensitive to GC7 than the parental HeLa S3 cells, indicating that MPV17L2 is not a GC7 target but has a synthetic lethal relationship with GC7 (Fig. 6B).

**Figure 6.**
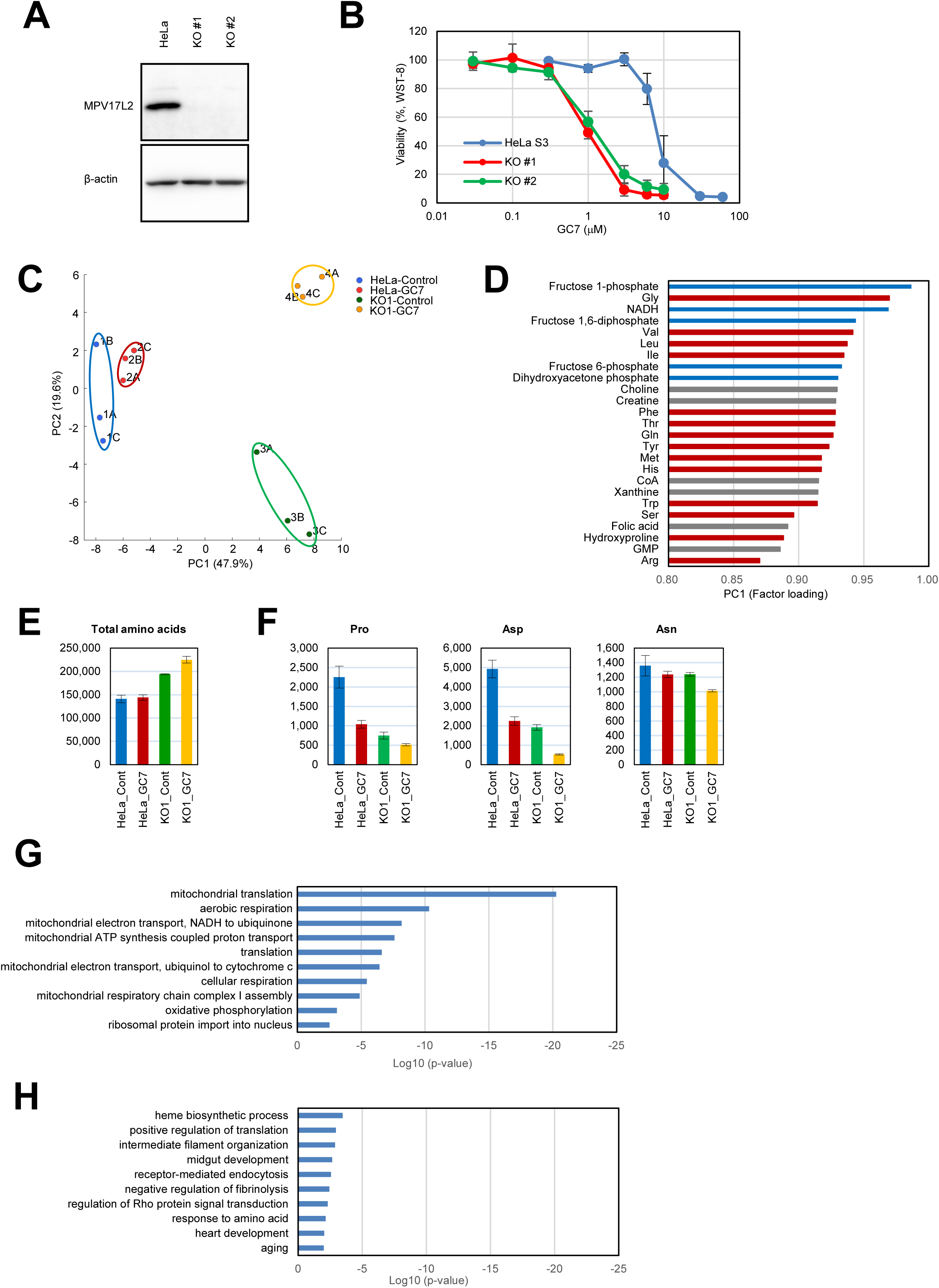
*MPV17L2* KO induces metabolic changes in cells. (A) Lysates from *MPV17L2* KO cells were analyzed by western blotting with anti-MPV17L2 antibody. (B) The sensitivity of HeLa S3 cells and *MPV17L2* KO cells to GC7 was analyzed by WST-8 assay (C–F) Metabolome analysis of HeLa S3 cells and KO #1 cells treated with 10 μM GC7 for 24 h. (C) Principal component analysis; (D) factor loading in PC1 (red bars for amino acids, blue for TCA cycle metabolites and gray for others); (E) amount of total amino acids; and (F) amount of Pro, Asp, and Asn. (G–H) Comparison of the proteome between HeLa S3 cells and *MPV17L2* KO cells. Proteins decreased (G) or increased (H) in *MPV17L2* KO #1 cells compared with HeLa S3 cells were subjected to pathway analysis.

To investigate the role of mitochondrial function, we performed a metabolome analysis of *MPV17L2* KO clone #1 and the parental HeLa S3 cells treated with and without 10 μM GC7 for 24 h. In the principal component analysis, the first principal component (PC1) clearly separated the parental cells and the KO cells, whereas PC2 separated GC7-treated and untreated cells (Fig. 6C). Metabolites that had large positive factor loadings in PC1, including the glycolysis metabolites fructose 1-phosphate, fructose 1,6-diphosphate, fructose 6-phosphate, and dihydroxyacetone phosphate, as well as many amino acids, increased in KO cells (Fig. S4, S5, and 6D). The increase in a subset of amino acids in KO cells affected the total amino acid content (Fig. 6E). Arg, citrulline, ornithine, and putrescine, all of which are upstream of spermidine biosynthesis, showed a relatively large positive factor loading in PC2 and were increased by GC7 treatment (Fig. S5 and S6). Despite the general increase in amino acid content caused by GC7, Pro and Asp decreased in GC7-treated control cells, and the decrease was greater in *MPV17L2* KO cells (Fig. 6F). Asn followed a similar pattern, although the changes were less pronounced (Fig. 6F). The decrease in the levels of Pro and Asp was observed previously in response to related compounds, such as inhibitors of the electron transport chain and ATP synthase including oligomycin A, rotenone, antimycin, and myxothiazol.^71^ Supplementation of Pro and Asp in the culture medium moderately but reproducibly restored the GC7 sensitivity in KO cells (Fig. S5), suggesting that the availability of these amino acids plays an important role in the cytotoxicity of GC7.

To explore the mechanism by which MPV17L2 affects the GC7 sensitivity of cells, we performed proteome analysis. In this set of experiments, we identified 6017 proteins including 766 mitochondrial proteins (Fig. S7). As observed in parental cells, GC7 treatment for 24 h in *MPV17L2* KO cells downregulated many mitochondrial proteins including mitochondrial ribosomal proteins and respiratory chain components (Fig. S7).

Analysis of the proteome in the absence of GC7 showed that approximately 500 proteins were decreased or increased by more than 2-fold in KO cells compared with parental HeLa cells (Fig. S7). Proteins involved in mitochondrial translation and aerobic respiration were enriched in the downregulated group (Fig. 6G), whereas no specific pathways were associated with proteins upregulated in KO #1 cells (Fig. 6H). This analysis indicated that *MPV17L2* KO cells contained a subset of downregulated mitochondrial proteins.

The results of proteome analysis were confirmed by examining the levels of mitochondrial proteins in *MPV17L2* KO cells by western blotting (Fig. 7A). The subunits of respiratory chain complexes I–V were significantly decreased in KO cells (Fig. 7A). Consistent with these results, mitochondrial networks were disturbed in KO cells (Fig. 7B). These phenotypes were enhanced by GC7 treatment (Fig. 7A and 7B).

**Figure 7.**
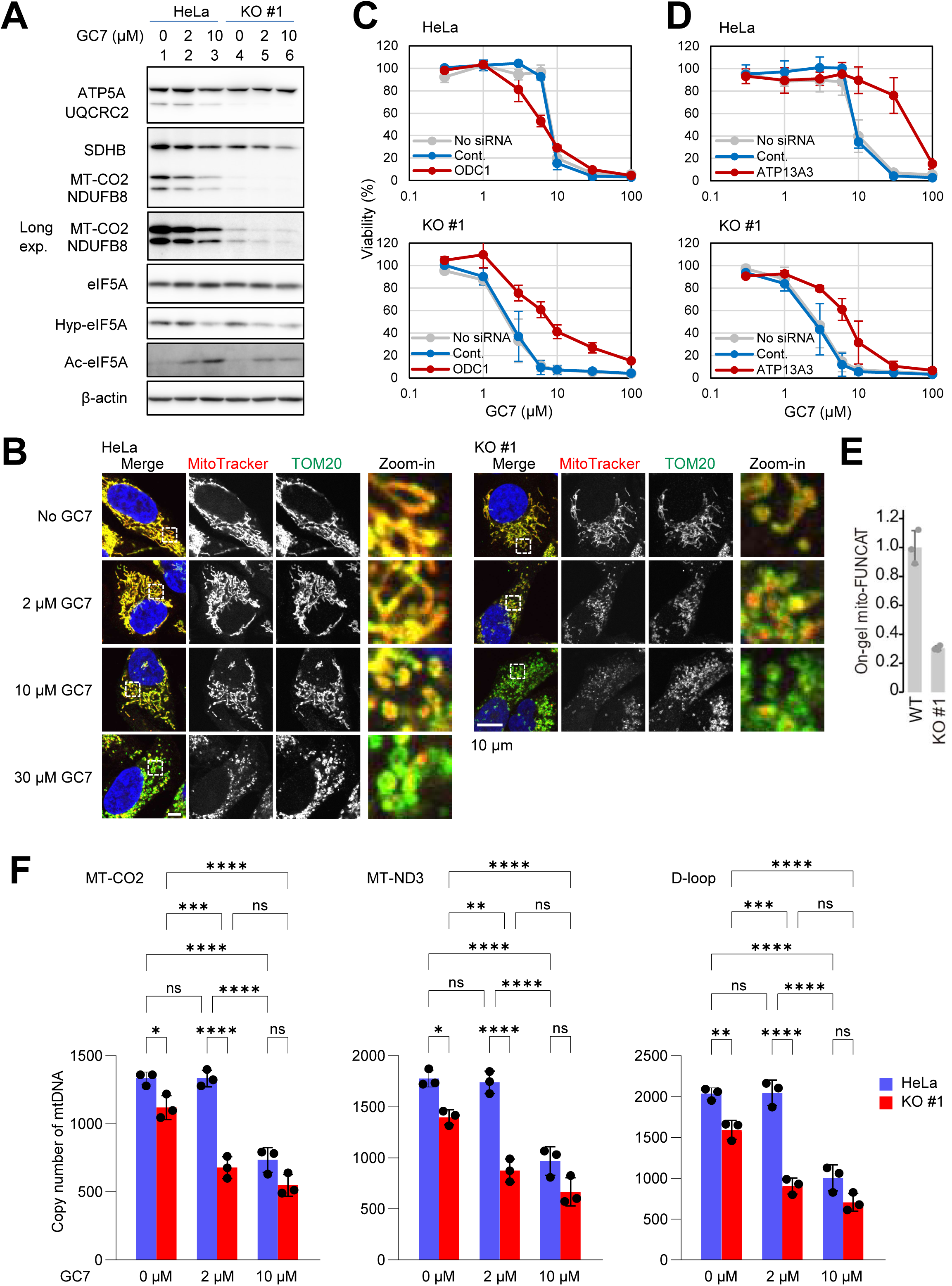
*MPV17L2* KO cells show aberrant mitochondrial morphology and decreased levels of mtDNA. (A) Lysates from HeLa S3 and KO #1 cells were analyzed by western blotting. (B) Mitochondrial morphology in HeLa S3 and KO #1 cells treated with GC7 for 24 h. (C) Effect of ODC1 siRNA knockdown on the sensitivity to GC7 in HeLa S3 (top) and KO #1 (bottom) cells was analyzed by WST-8 assay. Cont., non-targeting siRNA. (D) Effect of ATP13A3 siRNA knockdown on the sensitivity to GC7 in HeLa S3 (top) and KO #1 (bottom) cells was analyzed by WST-8 assay. (E) Mitochondrial translation activity of HeLa S3 and KO #1 cells was analyzed by in-gel mito-FUNCAT assay. (F) The mtDNA from HeLa S3 (blue) and KO #1 (red) cells treated with GC7 was quantified by qPCR. Data are expressed as the mean ± S.D. \**P* < 0.05, \*\**P* < 0.01, \*\*\**P* < 0.001, \*\*\*\**P* < 0.0001 (one-way ANOVA).

The potential genetic interaction between MPV17L2 and genes involved in GC7 sensitivity was examined by knockdown experiments using siRNAs. Knockdown of ODC1, which depleted putrescine from cells (Fig. S6), sensitized parental cells to GC7, whereas in *MPV17L2* KO cells, it decreased the sensitivity to GC7, suggesting genetic suppression of *MPV17L2* loss by ODC1 knockdown (Fig. 7C). By contrast, ATP13A3 knockdown caused both parental cells and *MPV17L2* KO cells to become less sensitive to GC7 (Fig. 7D), suggesting that there is no genetic interaction between *MPV17L2 and ATP13A3.*

The decrease in mitochondrial ribosomal proteins suggested that MPV17L2 depletion attenuated translation within the organelle. We used mitochondrial-specific fluorescent noncanonical amino acid tagging (FUNCAT) (mito-FUNCAT), in which newly synthesized polypeptides in mitochondria are pulse-labeled with the methionine analogue L-homopropargylglycine (HPG), followed by conjugation to fluorophores via the click reaction.^72^ Inhibition of cytosolic translation by anisomycin allowed the detection of translation in mitochondria specifically. The results showed that *MPV17L2* KO cells possessed low mitochondrial translation activity (∼30% of that in control cells) (Fig. 7E).

The decrease in mtDNA induced by GC7 treatment in HeLa cells (Fig. 2E) prompted us to examine the amount of mtDNA in *MPV17L2* KO cells (Fig. 7F). The results showed that the mtDNA content was lower in *MPV17L2* KO cells than in control cells. Treatment with a low dose of GC7 (2 μM) further decreased the mtDNA in KO cells significantly, but had no effect on mtDNA in control cells. Treatment with a high dose of GC7 (10 μM) decreased the mtDNA content in both cell groups.

Taken together, the results show that depletion of MPV17L2 caused various mitochondrial dysfunctions, including mtDNA loss, mitochondrial protein downregulation, and mitochondrial translation impairment, resulting in a phenocopy of the effects of GC7. The combination of the loss of *MPV17L2* with GC7 enhanced the phenotype and led to synthetic lethality.

## Discussion

In this study, we showed that the DHS inhibitor GC7 acts as a mitochondrial stressor. GC7 treatment downregulated many mitochondrial proteins encoded by the nuclear genome. Hypusinated eIF5A promotes translation elongation and termination by preventing ribosome collision at unfavorable nucleotide sequences or codons.^9,15,16,73^ Inhibition of DHS is thus expected to affect the translation of a subset of mRNAs. The mitochondrial targeting sequences of mitochondrial proteins are responsible for the efficient translation of their encoding mRNAs in a manner dependent on eIF5A hypusination.^12^ In this study, GC7 treatment downregulated mitochondrial ribosomal proteins, mitochondrial RNA polymerase, and TFAM at the post-transcriptional level. Consistent with the role of TFAM in the maintenance of mtDNA integrity,^74,75^ we found that the level of mtDNA was decreased in HeLa cells treated with GC7. This may have been responsible, at least in part, for the significant repression of transcriptional activity, together with the decrease of mitochondrial RNA polymerase and, in turn, led to the decrease in mtDNA-encoded proteins.

Treatment with mitochondrial stress inducers downregulates mitochondrial ribosomal proteins and activates ATF4 and the amino acid biosynthesis pathway.^63,76–79^ Treatment with GC7 or depletion of eIF5A induces the ISR, including PERK activation, eIF2α phosphorylation, and activation of the ATF4 protein (Ref. ^48,64,80^ and this study). ATF4 then induces the transcription of genes involved in amino acid synthesis.^81,82^ Consistently, we found an increase in many amino acids, with important exceptions such as the decrease in Pro and Asp, which is in line with the role of eIF5A in translation elongation of difficult-to-translate motifs. A previous study showed that treatment with oligomycin A causes a decrease in Pro.^71^ In this study, treatment with GC7 as well as FCCP and oligomycin A induced mitochondrial fragmentation, which is indicative of mitochondrial stress. Although it remains unclear whether the effect of GC7 on mitochondria is mediated by another pathway in addition to the inhibition of eIF5A hypusination, the present results indicate that GC7 induces mitochondrial stress leading to the ISR.

The nucleus and mitochondria communicate with each other.^58,83,84^ The nucleus controls mitochondrial activity by modulating the expression of nuclear-encoded mitochondrial proteins.^57,85,86^ Conversely, mitochondrial dysfunction, including damage to mtDNA, loss of mitochondrial membrane potential, and impairment of mitochondrial proteostasis, all of which were detected in response to GC7 treatment in this work, signals to the nucleus and induces a stress response regulating cytosolic translation via the ISR and mTORC1 activity.^58,77,87–89^

In this study, shRNA library screening identified genes that affect the sensitivity to GC7. Among them, we found genes encoding enzymes and proteins that regulate cellular polyamine content. Knockdown of genes encoding enzymes involved in spermidine synthesis sensitized cells to GC7 compared with control siRNAs, whereas knockdown of the spermine synthase (SMS), which utilizes spermidine, made cells less sensitive to GC7. The effect of GC7 on cell proliferation is correlated with its effect on inhibiting DHS, as indicated by comparison with a variety of similar compounds.^33^ Although we cannot exclude the possibility that GC7 competes with the cellular function(s) of spermidine itself or enzyme(s) that utilizes spermidine other than DHS,^48,90^ our results provide evidence that the cytotoxicity of GC7 depends on spermidine availability in the metabolism of polyamines.

The screening identified the polyamine transporter ATP13A3, the knockdown of which decreased the sensitivity to GC7. In mammalian cells, the P-type ATPases ATP13A2^91^ and ATP13A3^66^ at endosomes transport polyamines as well as several solute carriers such as SLC3A2, SLC7A5, SLC22A2, and SLC18B1, which localize to the plasma membrane or vesicles.^92–96^ Spermidine is a competitive antagonist of GC7 uptake, and cells with mutations in the *ATP13A3* gene are also defective in GC7 uptake.^33,66^ Given the structural similarity between GC7 and spermidine, the present results are consistent with a model in which depletion of ATP13A3 prevents the import of GC7 by endocytosis, rendering cells less sensitive to GC7.

In addition to the polyamine metabolism genes mentioned above, the shRNA screen identified *MPV17L2* as a gene whose knockdown increased the sensitivity of cells to GC7. MPV17L2 is important for mitochondrial translation by maintaining the integrity of mitochondrial ribosomal subunits, which leads to deficiency in the respiratory chain complex I subunits.^70,97^ We found that both overexpression and depletion of MPV17L2 induced morphological changes in mitochondria, and its depletion led to a decrease in mitochondrial ribosomal proteins and mitochondrial translation activity. Decreased mitochondrial translation induces a metabolic shift from OXPHOS to glycolysis due to the failure in the synthesis of respiratory chain complexes.^98^ MPV17L2 is homologous to three related proteins, the mitochondrial proteins MPV17 and MPV17L and the peroxisomal protein PXMP2.^70^ Among them, mutations in the human *MPV17* gene are implicated in mitochondrial diseases due to mtDNA depletion.^68,69,99^ Similarly, *MPV17L* knockout in human hepatocellular carcinoma HepG2 cells decreased mtDNA and several mitochondrial proteins including TFAM.^100^ These results raise the possibility of a causal relationship between mutations in the *MPV17L2* gene and human mitochondrial diseases; however, to the best of our knowledge, this association has not been reported to date. *MPV17L2* gene knockout significantly reduced mtDNA copy number (Fig. 7F). In addition, the sensitivity to GC7 was inversely related to MPV17L2 protein expression in cells, and the decrease in mtDNA and proteins in *MPV17L2* KO cells occurred at a lower concentration of GC7 than that in wild-type cells. These results suggest a synthetic lethal interaction between MPV17L2 and the action of GC7 and provide further evidence of the relationship between mitochondrial homeostasis and eIF5A hypusination. The hypusination of eIF5A, which regulates cytosolic translation of mitochondria-localized proteins, and mitochondrial translation of mtDNA-encoded proteins, which is promoted by MPV17L2, cooperatively regulate mitochondrial proteostasis and integrity. The present results also provide insight into the relationship between the eIF5A hypusination and mitochondrial diseases.

## Methods

### Cell lines

The human cervical cancer cell line HeLa S3 was cultured as described previously.^62^ Cell viability was measured using the WST-8 assay as described previously.^62^ HeLa ρ0 cells were generated under growth conditions including ethidium bromide and maintained at 37°C and 5% CO_2_ in DMEM supplemented with 10% fetal bovine serum,

1 mM sodium pyruvate, 5 μg/ml uridine, 100 units/ml penicillin, and 100 μg/ml streptomycin. To measure cell viability in HeLa ρ0 cells, cells cultured in 96-well plates were fixed with 4% paraformaldehyde, stained with 5 μg/ml Hoechst 33342 (Thermo Fisher) in phosphate-buffered saline, and counted using the Celigo Image Cytometer (Nexcelom Bioscience).

To generate MPV17L2 overexpressing cell line OE #1 and the empty vector-transfected cell line, pCI-neo-MPV17L2-FLAG and the empty vector pCI-neo-FLAG were transfected into HeLa S3 cells, which were then subjected to limiting dilution cloning by G418 selection. These cell lines were cloned and analyzed by western blotting.

To generate *MPV17L2* knockout cell lines, CAG-T7-hspCas9-T2A-RFP-H1-MPV17L2-gRNA#1 and CAG-T7-hspCas9-T2A-RFP-H1-MPV17L2-gRNA#2 plasmids were transfected into HeLa S3 cells using Lipofectamine 3000 reagent (Thermo Fisher). RFP-expressing cells were concentrated by fluorescence-activated cell sorting and subjected to limiting dilution cloning. *MPV17L2* KO cell lines #1 and #2 were cloned from cells transfected with the knockout plasmids #1 and #2, respectively.

### Chemicals

GC7 (LGC Biosearch Technologies), oligomycin A (Sigma), and FCCP (Sigma) were purchased.

### Plasmid construction

To generate pCI-neo-FLAG, two oligonucleotides for a FLAG-tag (5′-GGCCTTGACTACAAGGACGACGATGACAAGTAGGC-3′ and 5′-GGCCGCCTACTTGTCATCGTCGTCCTTGTAGTCAA-3′) were phosphorylated at the 5′ ends, annealed, and inserted into NotI-digested pCI-neo (Promega). MPV17L2 cDNA was obtained from total RNA of HeLa S3 cells and cloned into the NheI and XbaI sites of pCI-neo-FLAG to yield pCI-neo-MPV17L2-FLAG.

To construct *MPV17L2* knockout plasmids CAG-T7-hspCas9-T2A-RFP-H1-MPV17L2-gRNA#1 and CAG-T7-hspCas9-T2A-RFP-H1-MPV17L2-gRNA#2, oligonucleotides containing guide RNA sequences for MPV17L2 (#1, 5′-CTACGCCGCCTGTTATCCGC-3′ and #2, 5′-AGCGCCGTGGGTCGAAAACC-3′), designed by CRISPRdirect,^101^ were cloned into linearized CAG-T7-hspCas9-T2A-RFP-H1-gRNA SmartNuclease vector (CAS721R-1; System Biosciences).

### Proteome analysis

HeLa S3 and *MPV17L2* KO #1 and #2 cells cultured in 6-well plates were treated with or without 10 μM GC7. Cells were dissolved in 7 M guanidine-HCl, 1 M Tris-HCl (pH 8.5), 10 mM EDTA, and 50 mM dithiothreitol. After alkylation with iodoacetic acid, samples were desalted using the PAGE Clean Up Kit (Nacalai Tesque Inc.). The resultant precipitates were dissolved in 20 mM Tris-HCl (pH 8.0), 0.03% (w/v) n-dodecyl-β-D-maltoside, and digested with trypsin (tosyl phenylalanyl chloromethyl ketone treated; Worthington Biochemical Co) at 37°C overnight. Three technical replicates of each peptide were subjected to LC-MS/MS analysis. Solvent A (0.1% formic acid) and solvent B (80% acetonitrile with 0.1% formic acid) were used as LC solvents. Peptides were separated using an Easy nLC 1200 (Thermo Fisher Scientific) equipped with a nano-ESI spray column (NTCC-360, 0.075 mm internal diameter × 150 mm length, 3 μm, Nikkyo Technos Co) at a flow rate of 300 nl/min under a linear gradient condition over 250 minutes and analyzed with a Q Exactive HF-X Mass Spectrometer (Thermo Fisher Scientific) using the data-dependent Top 10 method.

The acquired data were processed using MASCOT 2.6 or 2.8 (Matrix Science) and Proteome Discoverer 2.2 or 2.4 (Thermo Fisher Scientific). The MASCOT search was conducted as follows: database, NCBIprot; taxonomy, homo sapiens (human); type of search, MS/MS ion; enzyme, trypsin; fixed modification, carboxymethyl (C); variable modifications, acetyl (protein N-term), Gln->pyro-Glu (N-term Q), oxidation (M); mass values, monoisotopic; peptide mass tolerance, ±15 ppm; fragment mass tolerance, ±30 mmu; max missed cleavages, 3; instrument type, ESI-TRAP. The acquired proteomic data were compared with MitoCarta 3.0^102^, and analyzed using QIAGEN Ingenuity Pathway Analysis.^103^

### RNA-seq

HeLa S3 cells cultured in 6-well plates were treated with 10 μM GC7 or 10 mM acetic acid (vesicle) for 24 h. Total RNA was isolated with TRI reagent (Sigma) and the RNeasy mini kit (Qiagen). RNA-seq was performed at Hokkaido System Science (Sapporo, Japan). Briefly, poly(A)-tailed RNA was isolated using oligo(dT)-beads, and next generation sequencing libraries that were prepared with NEBNext Ultra II Directional RNA Library Prep Kit for Illumina were sequenced at paired-end 150 bp on an Illumina NovaSeq 6000.

### Western blotting

Preparation of cell lysates and western blotting were performed as described previously.^104,105^ The primary antibodies used were as follows: hypusinated-eIF5A and acetylated-eIF5A,^106^ OXPHOS cocktail, HADHA, mitofusin 2, and ATP5B from Abcam, MPV17L2 (HPA043111), FLAG and β-actin from Sigma, TFAM and OPA1 from Cell Signaling Technology, eIF5A from BD Biosciences, mitochondrial RNA polymerase (POLRMT) from Novus, and MRPS17 from Proteintech.

### Immunocytochemistry

To examine changes in mitochondrial morphology, cells were stained with Mitotracker Red CMXRos (Thermo Fisher) and then immunostained with anti-Tom20 antibody (Santa Cruz Biotechnology) as described previously.^107,108^ Cells were examined under a confocal microscope (Olympus FV3000) using the objective lens UPLSAPO100XO (NA 1.4) or UPLSAPO60XS (NA 1.42).

### Nascent RNA analysis and quantitative PCR

HeLa S3 cells cultured in 6-well plates were treated with 10 μM GC7 for 2, 8, and 24 h. At 1 h before cell harvest, 0.5 mM EU was added. RNA was prepared using the TRI reagent, and EU-labeled RNA was recovered using Click-iT Nascent RNA Capture Kit (Thermo Fisher). Synthesis of cDNA and quantitative PCR (qPCR) were performed as described previously.^62^ The level of each nascent RNA was normalized to GAPDH nascent RNA. qPCR primer sets are listed in Table S1.^62,109–111^

### Mitochondrial DNA analysis

Genomic DNA was isolated from cells using GenElute Mammalian Genomic DNA Miniprep Kit (Sigma). qPCR was performed as described above. mtDNA copy number relative to nuclear DNA was calculated as 2^ΔCt (Ct for mtDNA-coded gene – Ct for B2-macroglobulin).112^

### shRNA screening

shRNA screening was performed as described previously.^62,105,113^ Briefly, HeLa S3 cells were transduced with the Decipher Human Module 1; one aliquot of transduced cells was untreated and another aliquot was treated with 10 μM GC7 for 10 days (Exp. 1) or 14 days (Exp. 2) and then collected. The same experiments were repeated with Module 2-infected cells and Module 3-infected cells. Genomic DNA extraction, PCR amplification of barcodes with indexed primers, and barcode quantitation by next-generation sequencing on an Illumina HiSeq 2500 (Macrogen Japan) were performed as described previously.^105^ We selected genes with three or more shRNAs with a threshold of fold change (GC7-treated/untreated) <0.5 and >2 (Exp. 1) or <0.5, <0.33 and >2 (Exp. 2).

### siRNA knockdown

HeLa S3 cells were transfected with 10–20 nM siRNA pools (Human ON-TARGETplus SMARTpool, Dharmacon) containing four siRNAs against each gene, or with ON-TARGETplus Non-Targeting siRNA #1 using Lipofectamine RNAiMAX reagent (Thermo Fisher Scientific). After 72 h, transfected cells were split and seeded in 96-well plates, and treated with various concentrations of GC7 for 72 h.

### Metabolome analysis

HeLa S3 and *MPV17L2* KO #1 cells in 100 mm dishes were treated with or without 10 μM GC7 for 24 h. The cells were washed twice with 5% mannitol and extracted in 0.8 ml methanol and 0.55 ml distilled water containing 10 µM Internal Standard Solution (Human Metabolome Technologies, Tsuruoka, Japan). The extracts were cleared by centrifugation (2,300 × g for 5 min, 4°C), and centrifugally filtered through a Millipore 5-kDa cutoff filter to remove proteins. Analyses of the filtrate using CE-TOFMS were performed at Human Metabolome Technologies.

### On-gel mito-FUNCAT

On-gel mito-FUNCAT was performed as described previously.^72,114^ HeLa S3 and *MPV17L2* KO #1 cells were cultured in 6-well plates in methionine-free medium with 50 µM HPG (Jena Bioscience) and 100 µg/ml anisomycin (Alomone Labs, A-520) for 3 h before cell harvest. After lysis of cells, nascent peptides were labeled with 50 μM IRdye800CW Azide (LI-COR Biosciences) using a Click-it Cell Reaction Buffer Kit (Thermo Fisher Scientific) according to the manufacturer’s instructions. The supernatants were separated by SDS–PAGE, and the signals were detected using an Odyssey CLx (LI-COR Biosciences) with an IR 800-nm channel. The signals were standardized against Coomassie Brilliant Blue staining using the IR 700-nm channel.

### Ribosome profiling and disome profiling

Ribosome profiling and disome profiling library preparation was performed as previously reported.^8,115^ The data were analyzed as previously described.^8,116^

### Measurement of polyamines

HeLa S3 and *MPV17L2* KO #1 cells cultured in 6-well plates were transfected with ODC1 siRNA pool as described above. After 2 days, the culture was supplemented with GC7 and incubated for an additional 24 h before cell harvest. Cells were homogenized with 5% trichloroacetic acid, and the homogenates were incubated at 70°C for 10 min, followed by centrifugation at 17,000 *g* for 5 min. The pellets were used to determine the protein content using the DC protein assay (Bio-Rad). The supernatants were used to measure polyamine contents according to the method of Igarashi et al.^117^

### Quantification and statistical analysis

The statistical details can be found in figure legends. Statistical analyses were calculated using GraphPad Prism 10. One-way ANOVA was performed. These data are presented as the mean ± standard deviation (SD).

## Acknowledgements

We thank the members of Yoshida laboratory for helpful discussions and materials. The authors wish to thank N. Suzuki, M. Tagami, and S. Aoki (GeNAS, RIKEN) for help with deep sequencing and M. Muroi and H. Osada (RIKEN Center for Sustainable Resource Science) for help at the initial stage of this work. This work was supported in part by JSPS under Grants-in-Aid (21H02434 and 22H04922 to K.M.; 23H05473 and 23H04882 to M.Y.; and JP23H02415 to S.I.) from the Ministry of Education, Culture, Sports, Science and Technology of Japan. P.H. was supported by a JSPS (DC2) fellowship. T.W. was a recipient of fellowships from JSPS (DC2), JST SPRING (JPMJSP2108), and the ANRI.

