## Supplementary Information for "Chemical genetic interaction linking eIF5A hypusination and mitochondrial integrity"

### Table S1. Primer sequences used for qPCR.

| Gene | Forward primer (5' to 3') | Reverse primer (5' to 3') |
|---|---|---|
| MT-CO1 | GACGTAGACACACGAGCATATTTCA | AGGACATAGTGGAAGTGAGCTACAAC |
| MT-CO2 | CGTCTGAACTATCCTGCCCG | GAGGGATCGTTGACCTCGTC |
| MT-CO3 | ACCAATGATGGCGCGATGTA | GGCTGGAGTGGTAAAAGGCT |
| MT-ND1 | CTACTACAACCCTTCGCTGAC | GGATTGAGTAAACGGCTAGGC |
| MT-ND3 | GCGGCTTCGACCCTATATCC | AGGGCTCATGGTAGGGGTAA |
| MT-ND5 | TCGAATAATTCTTCTCACCC | TAGTAATGAGAAATCCTGCG |
| MT-ND6 | GTAGGATTGGTGCTGTGG | GGATCCTCCCGAATCAAC |
| MT-CYB | ATCACTCGAGACGTAAATTATGGCT | GGAGGATAATGCCGATGTTTCAG |
| MT-ATP6 | TAGCCATACACAACACTAAAGGACGA | GGGCATTTTTAATCTTAGAGCGAAA |
| MT-ATP8 | TACCACCTACCTCCCTCACC | AGGATTGTGGGGGCAATGAAT |
| RNR1 | CATCAAGCACGCAGCAATGCAG | GTTAATCACTGCTGTTTCCCGTG |
| RNR2 | CCAAGCATAATATAGCAAGGAC | CTTAGCTTTGGCTCTCCTTG |
| D-loop | CATCTGGTTCCTACTTCAGGG | CCGTGAGTGGTTAATAGGGTG |
| ATP5A | ATGACGACTTATCCAAACAGGC | CGGGAGTGTAGGTAGAACACAT |
| UQCRC2 | TAAGTGTGACCGCAACAAGGG | TGGTGACATTGAGCAGGAACT |
| SDHB | ACCTTCCGAAGATCATGCAGA | GTGCAAGCTAGAGTGTTGCCT |
| NDUFB8 | ACAGGAACCGTGTGGATACAT | CCCCACCCAGCACATGAAT |
| MPV17L2 | TTTCGAGTCACCTACATCAACG | ACTGGGCTCCGGTACTTCAA |
| GAPDH | GCACCGTCAAGGCTGAGAAC | TGGTGAAGACGCCAGTGGA |
| B2-macroglobulin | TGCTGTCTCCATGTTTGATGTATCT | TCTCTGCTCCCCACCTCTAAGT |
| MT-ND2 | Bio-Rad PrimePCR SYBR Green Assay: MT-ND2, Human Unique Assay ID: qHsaCED0034386 | |
| MT-ND4 | Bio-Rad PrimePCR SYBR Green Assay: MT-ND4, Human Unique Assay ID: qHsaCED0007715 | |
| MT-ND4L | Bio-Rad PrimePCR SYBR Green Assay: MT-ND4L, Human Unique Assay ID: qHsaCED0007738 | |

### Supplementary Figures and Legends

**Figure S1. Proteome and transcriptome analysis of GC7-treated HeLa S3 cells.**

(A) Proteins from HeLa S3 cells treated with 10 μM GC7 for 24 h or 72 h were analyzed by label-free mass spectrometry. Proteins commonly detected were compared with Mitocarta 3.0.

(B and C) MA plot and a heat map of transcriptome analysis using HeLa S3 cells treated with 10 μM GC7 for 24 h.

**Figure S2. Effect of GC7 on mitochondrially-encoded RNAs and mitochondrial morphology.**

(A) HeLa S3 cells were treated with GC7 for 24 h. Mitochondrially-encoded RNAs and two nuclear-encoded mRNAs from the cells were analyzed by qPCR.

(B) HeLa S3 cells were treated with GC7 for 1, 4, and 24 h. The cells were stained with Mitotracker Red and analyzed by immunofluorescence with anti-Tom20 antibodies. The cells were counter-stained with DAPI and observed by confocal microscopy.

**Figure S3. Establishment of MPV17L2 KO cells.**

(A) Gene editing was confirmed by amplicon deep sequencing in HeLa, KO #1, and KO #2 cells. Sequence ID (Seq_ID) s000001 was matched to region 18327314–18327708 of GenBank CP068259. The length of deletion or insertion of each amplicon is shown in the table. Percentage represents the population of each amplicon in total read counts.

(B) Alignment of amplicons is shown. Deletion and insertion of each amplicon, positions of the start codon, and gRNAs #1 and #2 are indicated in color. Seq_ID s000006 has an in-frame termination codon (underlined), although the length of insertion is a multiple of 3, indicating that only a short peptide is produced from this allele.

**Figure S4. GC7 treatment and MPV17L2 knockout both affected the amount of metabolites from glycolysis and the TCA cycle.**

Metabolome analysis of HeLa S3 cells and KO #1 cells treated with 10 μM GC7 for 24 h. Metabolites in glycolysis (A) and the TCA cycle (B) are shown.

**Figure S5. GC7 treatment and MPV17L2 knockout increased the amounts of amino acids except Pro, Asp, and Asn.**

(A) Amounts of 20 amino acids from the metabolome analysis.

(B) Factor loadings in PC2 (bars in red for amino acids, blue for TCA cycle metabolites, and yellow for polyamine pathway).

(C) The effect of the addition of 100 μM of each Pro and Asp (P + D) or a non-essential amino acid mixture (NEAA) on GC7 sensitivity in HeLa S3, KO #1, and KO #2 cells was analyzed by the WST-8 assay.

**Figure S6. Amounts of polyamines in HeLa S3 cells and KO #1 cells.**

(A) The amounts of ornithine and polyamines from the metabolome analysis are shown.

(B) Polyamines were quantified from HeLa S3 cells and KO #1 cells treated with 10 μM GC7 for 24 h after knockdown of ODC1. Data represent the mean values of two independent experiments. N.D., not detected.

**Figure S7. Proteome analysis of HeLa S3, KO #1, and KO #2 cells.**

HeLa S3, KO #1, and KO #2 cells were treated with 10 μM GC7 for 24 h. Cellular proteins were analyzed by label-free mass spectrometry. Proteins decreased (<0.5) or increased (>2) in cells treated with GC7 compared with untreated cells were analyzed against the list of mitochondrial proteins (MitoCarta 3.0) (A). Proteins decreased (<0.5) or increased (>2) in KO cells compared with HeLa S3 cells were analyzed against the list of mitochondrial proteins (B).

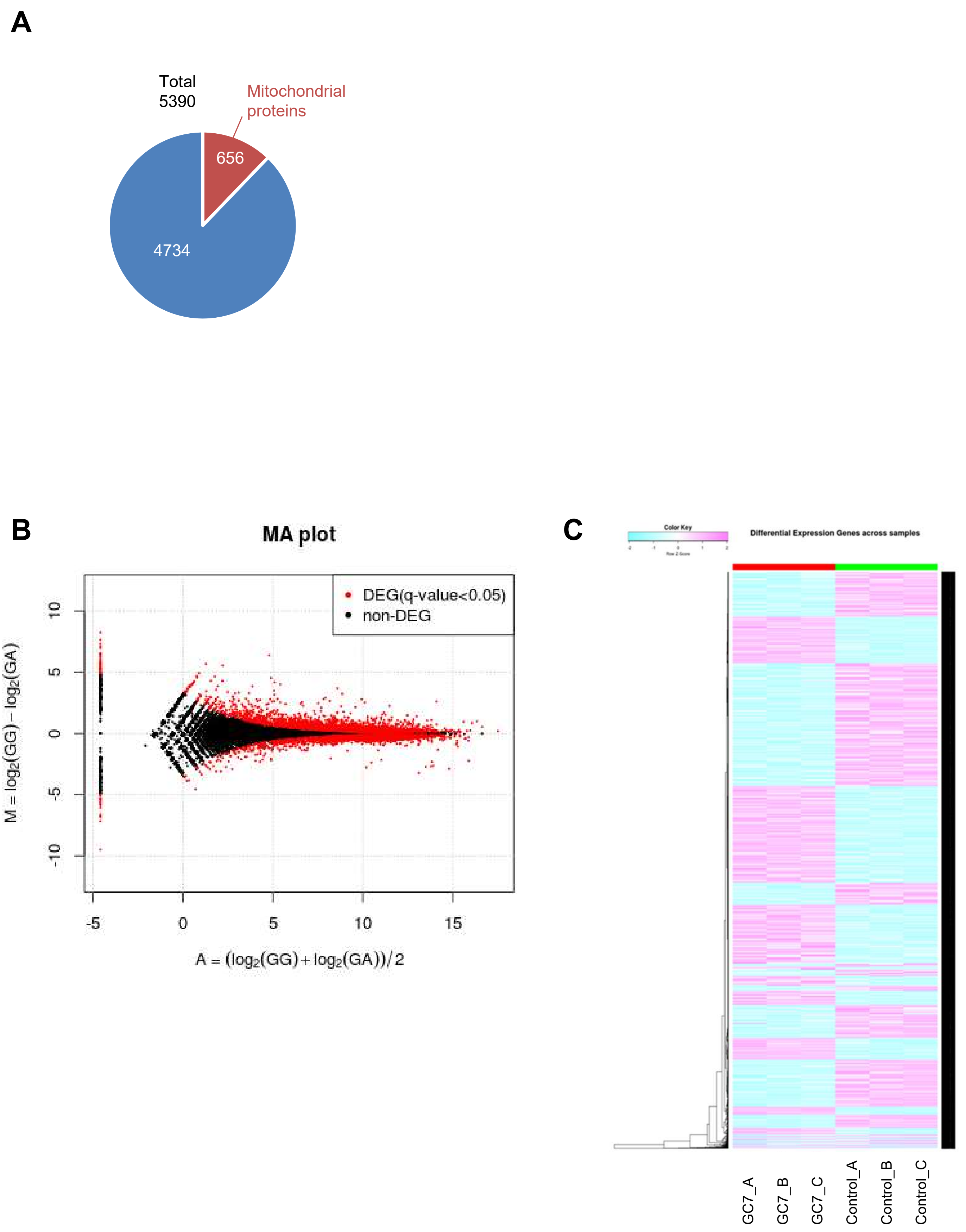


**Figure S1. Proteome and transcriptome analysis of GC7-treated HeLa S3 cells.**

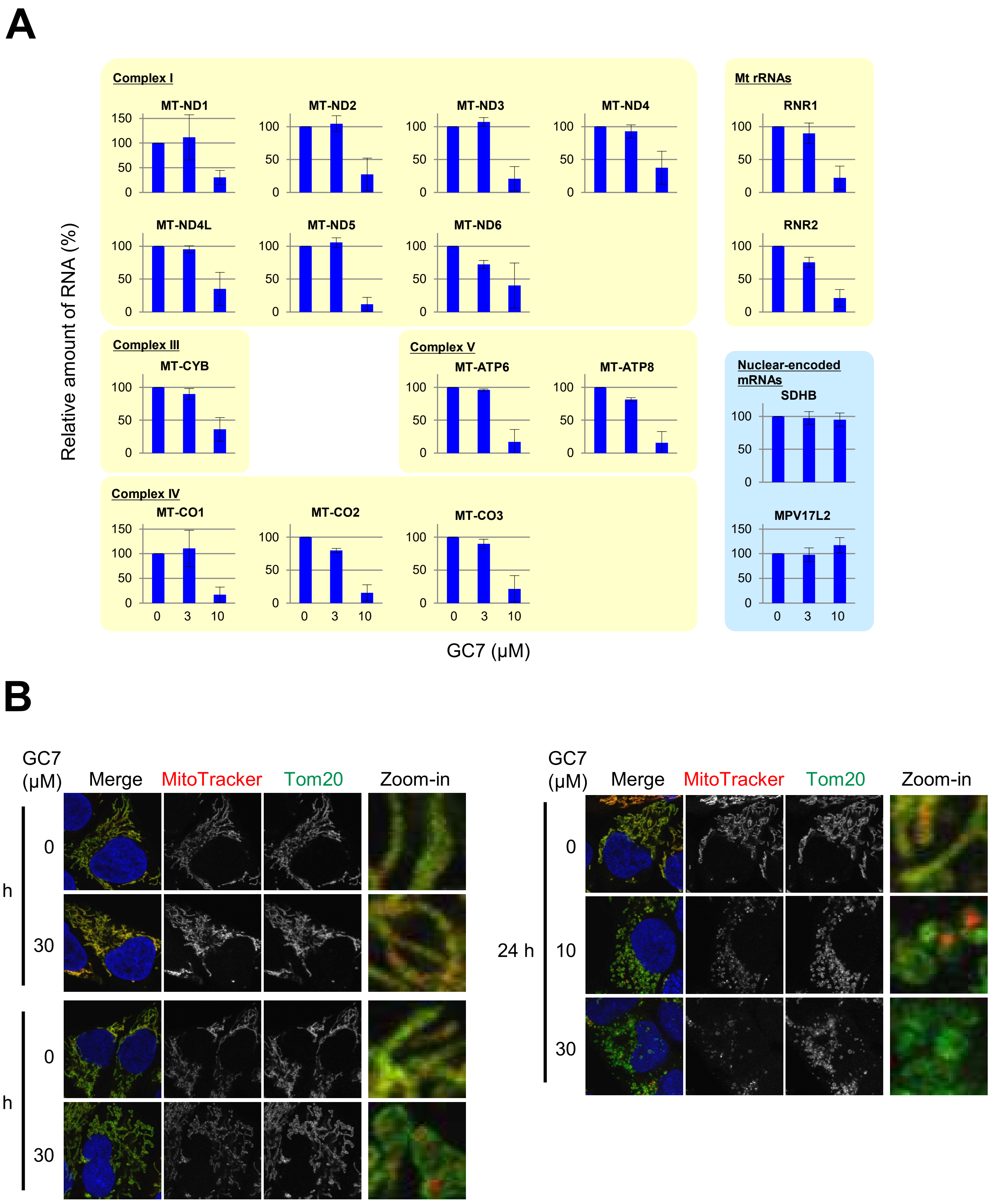


**Figure S2. Effect of GC7 on mitochondrially-encoded RNAs and mitochondrial morphology.**

**A**

| Seq_ID | | HeLa | KO #1 | KO #2 |
|---|---|---|---|---|
| s000001 | WT | 99.9% | 0.9% | 22.8% |
| s000002 | 205 del | 0.0% | 55.1% | 0.0% |
| s000003 | 1 del | 0.0% | 0.0% | 44.7% |
| s000004 | 68 del | 0.0% | 0.0% | 32.4% |
| s000005 | 1 del | 0.0% | 22.8% | 0.0% |
| s000006 | 27 insert | 0.0% | 21.2% | 0.0% |

**B**

```
                     *         *         *         *         *         *         *         *         *         *
s000001   1>AGGGGCGGGCAATTTGGGATCGACAGTGACTCGCGACTGGTCGGCGCGGCGAAAGCAGAGCGGCGCGCCGGTTCCTTGGTTCCTGAGGGCGATGGCGCGG>100
s000002   1>AGGGGCGGGCAATTTGGGATCGACAGTGACTCGCGACTGGTCGGCGCGGCGAAAGCAGAGCGGCGCGCCGGTTCCTTGGTTCCTGAGGGCGATGGCGCGG>100
s000003   1>AGGGGCGGGCAATTTGGGATCGACAGTGACTCGCGACTGGTCGGCGCGGCGAAAGCAGAGCGGCGCGCCGGTTCCTTGGTTCCTGAGGGCGATGGCGCGG>100
s000004   1>AGGGGCGGGCAATTTGGGATCGACAGTGACTCGCGACTGGTCGGCGCGGCGAAAGCAGAGCGGCGCGCCGGTTCCTTGGTTCCTGAGGGCGATGGCGCGG>100
s000005   1>AGGGGCGGGCAATTTGGGATCGACAGTGACTCGCGACTGGTCGGCGCGGCGAAAGCAGAGCGGCGCGCCGGTTCCTTGGTTCCTGAGGGCGATGGCGCGG>100
s000006   1>AGGGGCGGGCAATTTGGGATCGACAGTGACTCGCGACTGGTCGGCGCGGCGAAAGCAGAGCGGCGCGCCGGTTCCTTGGTTCCTGAGGGCGATGGCGCGG>100

             *         *         *                                    *         *         *         *
101>GGTGGCTGGCGCCGGCTACGCCGCCTGTTATC---------------------------CGCGGGGCAGCTTCTATTCCAGGGCCGCGCGCTGCTCGTCA>173
101>GGTGGCTGGC------------------------------------------------------------------------------------------>110
101>GGTGGCTGGCGCCGGCTACGCCGCCTGTTATC---------------------------CGCGGGGCAGCTTCTATTCCAGGGCCGCGCGCTGCTCGTCA>173
101>GGTGGCTGGCGCCGGCTACGCCGCCTGTTATC---------------------------CGCGGGGCAGCTTCTATTCCAGGGCCGCGCGCTGCTCGTCA>173
101>GGTGGCTGGCGCCGGCTACGCCGCCTGTTATC----------------------------GCGGGGCAGCTTCTATTCCAGGGCCGCGCGCTGCTCGTCA>172
101>GGTGGCTGGCGCCGGCTACGCCGCCTGTTATCGTTGATTCTTTGACCTAATTTAGTGAGCGCGGGGCAGCTTCTATTCCAGGGCCGCGCGCTGCTCGTCA>200
          *         *         *         *         *         *         *         *         *         *
174>CTAACACGCTGGGCTGCGGCGCGCTCATGGCGGCCGGTGATGGCGTGCGCCAGTCCTGGGAGATCCGCGCCCGGCCCGGCCAGGTTTTCGACCCACGGCG>273
110>---------------------------------------------------------------------------------------------------->110
174>CTAACACGCTGGGCTGCGGCGCGCTCATGGCGGCCGGTGATGGCGTGCGCCAGTCCTGGGAGATCCGCGCCCGGCCCGGCCAGGTTT-CGACCCACGGCG>272
174>CTAACACGCTGGGCTGCGGCGCGCTCATG--------------------------------------------------------------------GCG>205
173>CTAACACGCTGGGCTGCGGCGCGCTCATGGCGGCCGGTGATGGCGTGCGCCAGTCCTGGGAGATCCGCGCCCGGCCCGGCCAGGTTTTCGACCCACGGCG>272
201>CTAACACGCTGGGCTGCGGCGCGCTCATGGCGGCCGGTGATGGCGTGCGCCAGTCCTGGGAGATCCGCGCCCGGCCCGGCCAGGTTTTCGACCCACGGCG>300
          *         *         *         *         *         *         *         *         *         *
274>CTCCGGTGAGGACGCCACGCTGCTTAGTCCTTCACCCCGGGCGACCTTTGATCCCGACACCCGACTGCGGGCCGTGACCCCTGACCCGTGGCCTCAGTCC>373
111>------------------------------------------GACCTTTGATCCCGACACCCGACTGCGGGCCGTGACCCCTGACCCGTGGCCTCAGTCC>168
273>CTCCGGTGAGGACGCCACGCTGCTTAGTCCTTCACCCCGGGCGACCTTTGATCCCGACACCCGACTGCGGGCCGTGACCCCTGACCCGTGGCCTCAGTCC>372
206>CTCCGGTGAGGACGCCACGCTGCTTAGTCCTTCACCCCGGGCGACCTTTGATCCCGACACCCGACTGCGGGCCGTGACCCCTGACCCGTGGCCTCAGTCC>305
273>CTCCGGTGAGGACGCCACGCTGCTTAGTCCTTCACCCCGGGCGACCTTTGATCCCGACACCCGACTGCGGGCCGTGACCCCTGACCCGTGGCCTCAGTCC>372
301>CTCCGGTGAGGACGCCACGCTGCTTAGTCCTTCACCCCGGGCGACCTTTGATCCCGACACCCGACTGCGGGCCGTGACCCCTGACCCGTGGCCTCAGTCC>400
          *         *
374>CCCGCAGGACCCTTGCCCTGAC>395
169>CCCGCAGGACCCTTGCCCTGAC>190
373>CCCGCAGGACCCTTGCCCTGAC>394
306>CCCGCAGGACCCTTGCCCTGAC>327
373>CCCGCAGGACCCTTGCCCTGAC>394
401>CCCGCAGGACCCTTGCCCTGAC>422
```

Dark Green: gRNA1
Green: gRNA2 (complementary strand)
Blue: Start codon
Red: Deletion
Yellow: Insertion with a termination codon

**Figure S3. Establishment of MPV17L2 KO cells.**

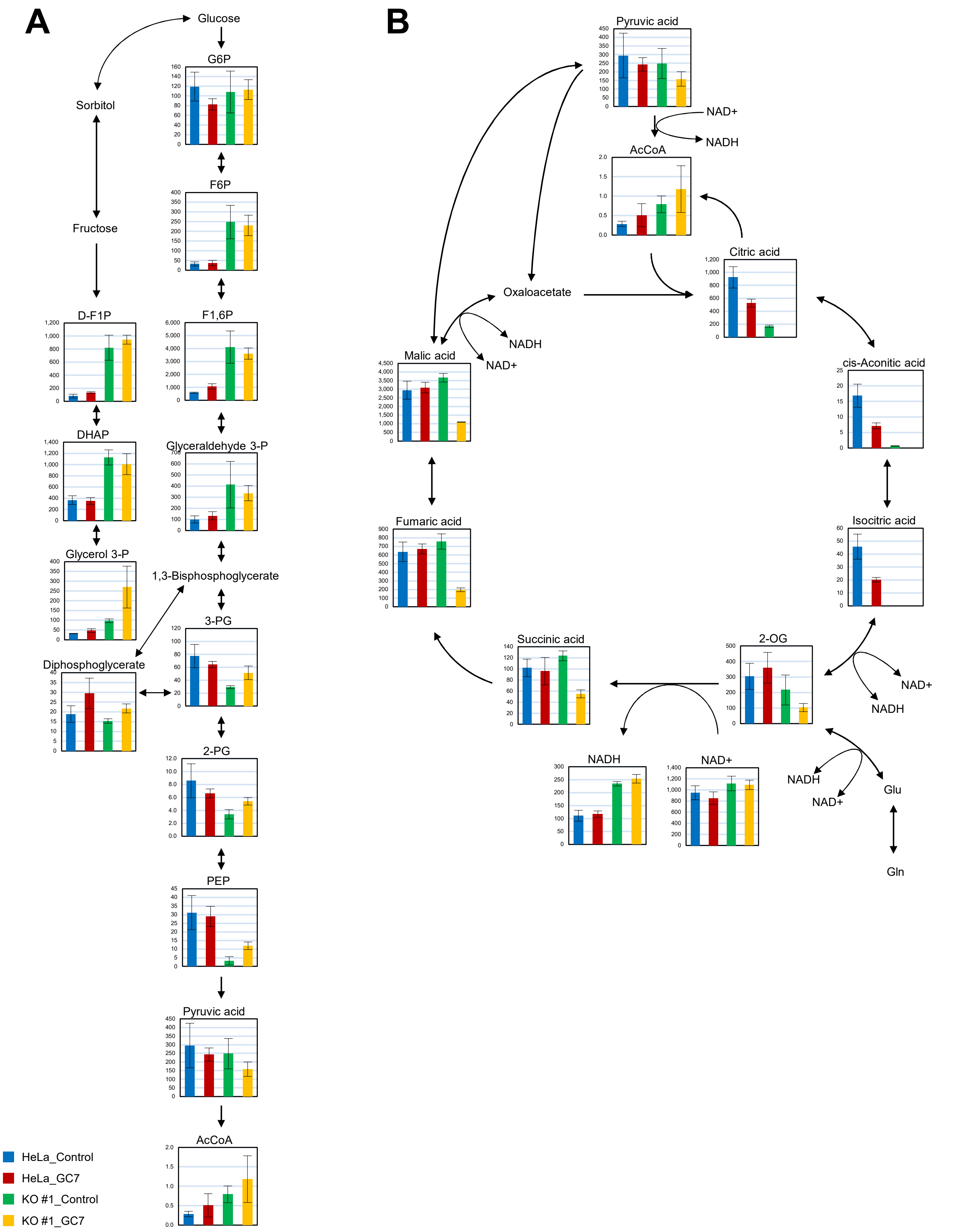


**Figure S4. GC7 treatment and MPV17L2 knockout both affected the amount of metabolites from glycolysis and the TCA cycle.**

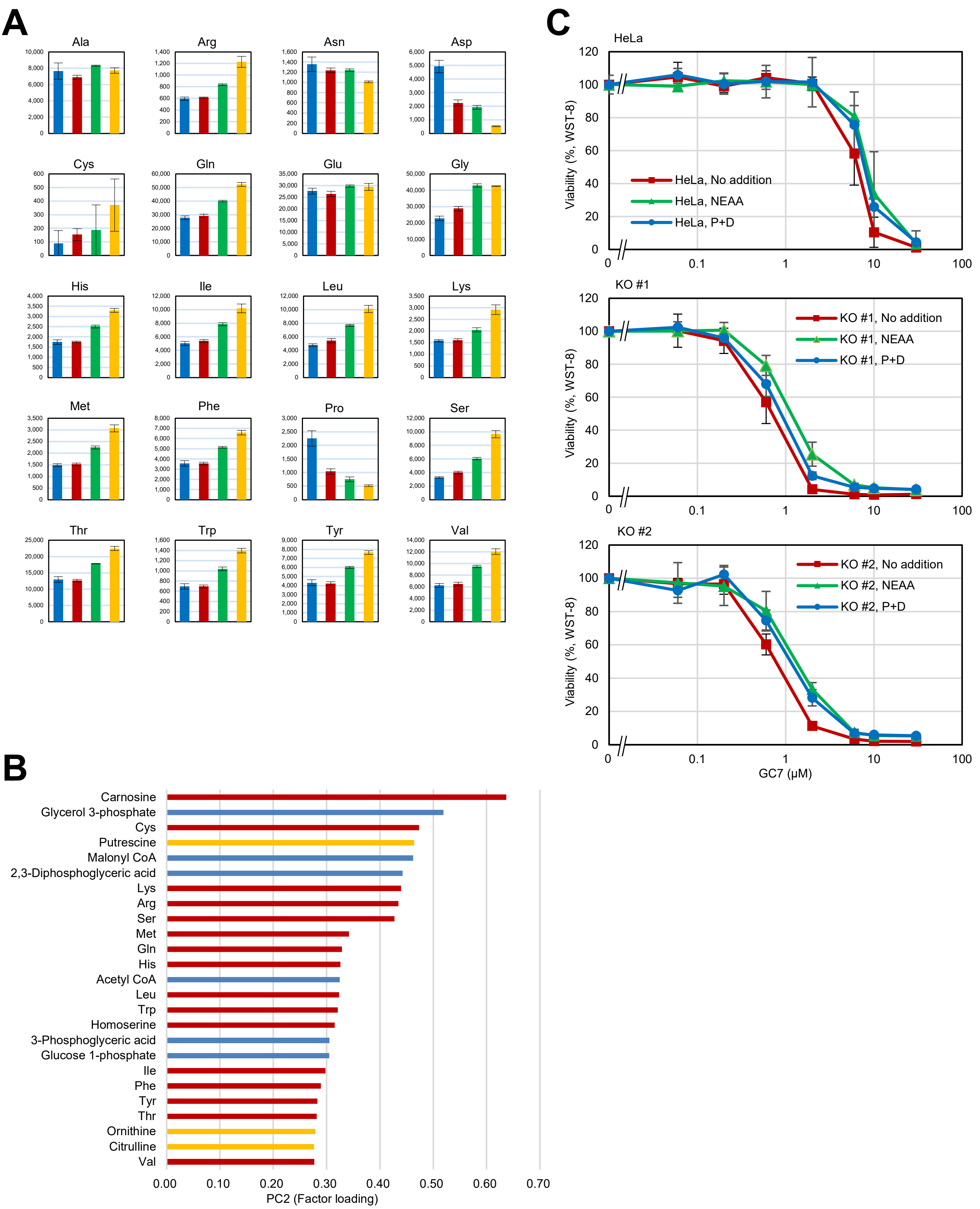


**Figure S5. GC7 treatment and MPV17L2 knockout increased the amounts of amino acids except Pro, Asp, and Asn.**

## A

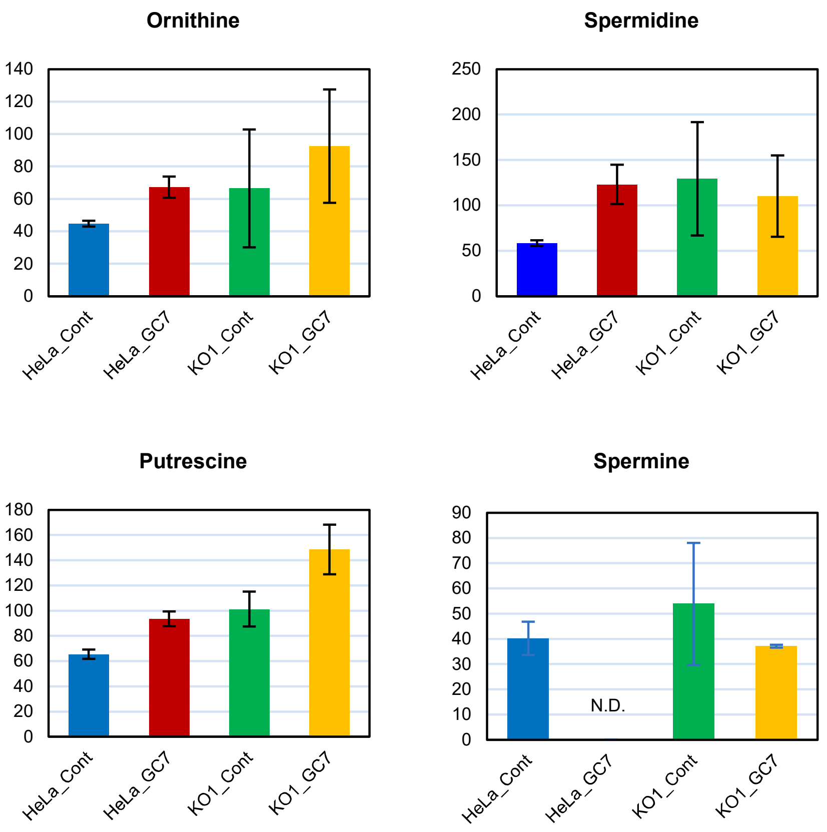


## B

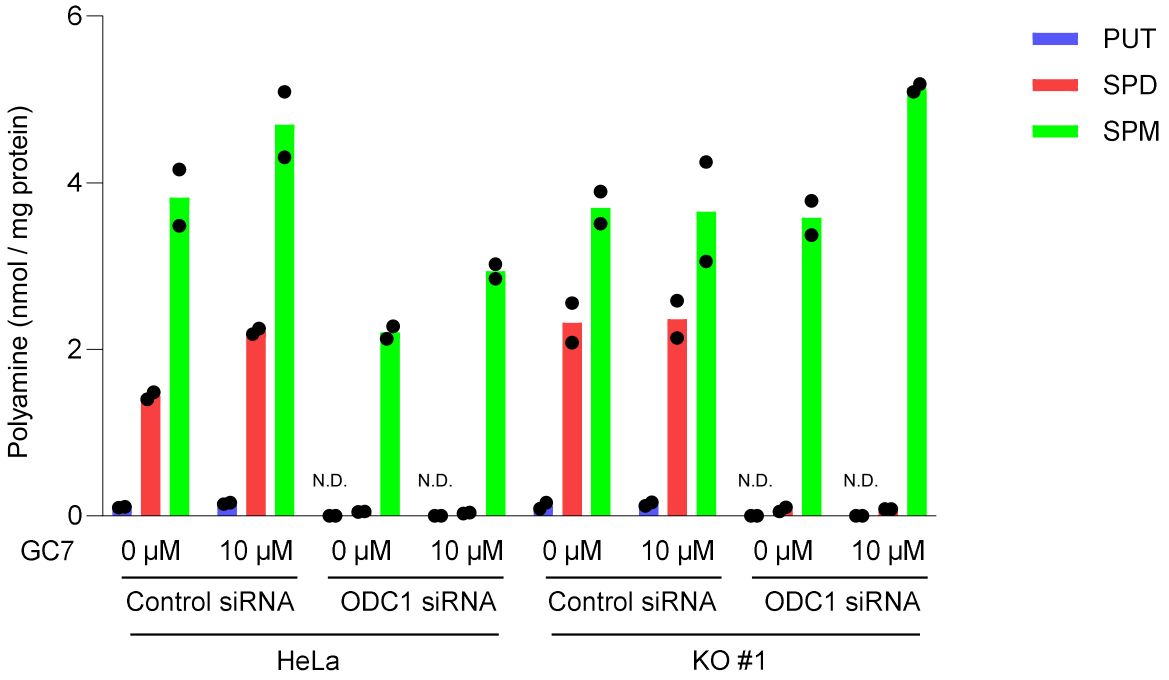


**Figure S6. Amounts of polyamines in HeLa S3 cells and KO #1 cells.**

A

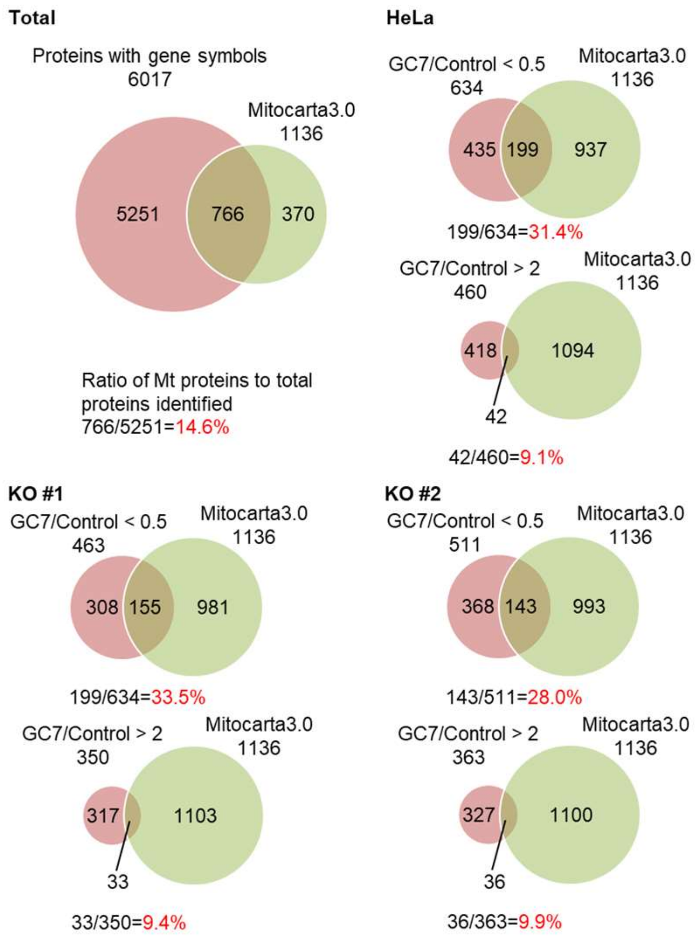


B

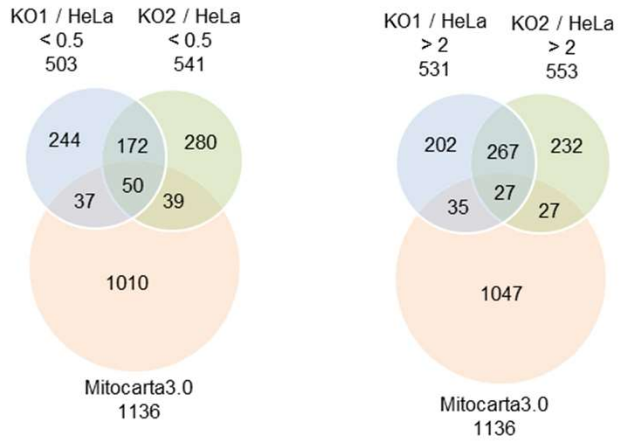


**Figure S7. Proteome analysis of HeLa S3, KO #1, and KO #2 cells.**
